# Autophagy preferentially degrades non-fibrillar polyQ aggregates

**DOI:** 10.1101/2023.08.08.552291

**Authors:** Dorothy Y. Zhao, Felix J.B. Bäuerlein, Itika Saha, F. Ulrich Hartl, Wolfgang Baumeister, Florian Wilfling

**Affiliations:** Max Planck Institute of Biochemistry, Molecular Machines and Signaling, 82152 Martinsried, Germany; Max Planck Institute of Biochemistry, Molecular Structural Biology, 82152 Martinsried, Germany; University Medical Center Göttingen, Institute of Neuropathology, 37099 Göttingen, Germany; Max Planck Institute of Biochemistry, Cellular Biochemistry, 82152 Martinsried, Germany; Max Planck Institute of Biophysics, Mechanisms of Cellular Quality Control, 60439 Frankfurt a. M., Germany; Aligning Science Across Parkinson’s (ASAP) Collaborative Research Network, Chevy Chase, MD 20815

**Keywords:** protein aggregation, neurodegeneration, polyglutamine/polyQ expansion, amyloid fibril, phase separation, autophagy, aggrephagy, p62/SQSTM1/sequestosome 1, cryo-electron tomography, cryo-electron microscopy

## Abstract

Aggregation of proteins containing expanded polyglutamine (polyQ) repeats is the cytopathologic hallmark of a group of dominantly inherited neurodegenerative diseases, including Huntington’s disease (HD). Huntingtin (Htt), the disease protein of HD, forms amyloid-like fibrils by liquid-to-solid phase transition. Macroautophagy has been proposed to clear polyQ aggregates, but the efficiency of aggrephagy is limited. Here, we used cryo-electron tomography to visualize the interactions of autophagosomes with polyQ aggregates in cultured cells *in situ*. We found that an amorphous aggregate phase exists next to the radially organized polyQ fibrils. Autophagosomes preferentially engulfed this amorphous material, mediated by interactions between the autophagy receptor p62/SQSTM1 and the non-fibrillar aggregate surface. In contrast, amyloid fibrils excluded p62 and evaded clearance, resulting in trapping of autophagic structures. These results suggest that the limited efficiency of autophagy in clearing polyQ aggregates is due to the inability of autophagosomes to interact productively with the non-deformable, fibrillar disease aggregates.

## Introduction

Numerous neurodegenerative disorders (ND) including Alzheimer’s disease, Parkinson’s disease, amyotrophic lateral sclerosis and Huntington’s disease, are associated with the formation of toxic aggregates in neuronal cells (Hipp et al., 2019; Ross and Poirier, 2004; Taylor et al., 2002). Huntington’s disease, characterized by a progressive motor and cognitive decline, is the most frequent member of a group of diseases caused by dominantly-inherited polyglutamine (polyQ) repeat expansions in otherwise unrelated proteins (Gusella and MacDonald, 2006; Walker, 2007; Williams and Paulson, 2008). Expansion of the polyQ sequence in exon 1 of the protein huntingtin (Htt) beyond ∼37Q is a strong predictor of Huntington’s disease (Duyao et al., 1993; Group, 1993; Gusella and MacDonald, 2006; Walker, 2007). The length of the polyQ tract correlates positively with aggregation propensity and disease severity and inversely with the age of disease onset (Gusella and MacDonald, 2006; Walker, 2007). Longer repeats (up to >100 Q) more readily form amyloid-like fibrils, giving rise to highly stable nuclear and cytosolic inclusion bodies (DiFiglia et al., 1997; Scherzinger et al., 1997), initially in striatal neurons and then also in other brain regions as pathology progresses (DiFiglia et al., 1997; Gusella and MacDonald, 2006; Walker, 2007).

Aggregates of polyQ-expanded Htt exhibit structurally different forms, including dynamic soluble oligomers and stable fibrils with cross-β structure (Bauerlein et al., 2020; Bauerlein et al., 2017; Chiti and Dobson, 2017; Gruber et al., 2018; Hoop et al., 2016; Kim et al., 2016; Miller et al., 2011; Nucifora et al., 2012; Scherzinger et al., 1997). Recently, an amorphous gel-like state has been described as an intermediate stage of fibril formation in yeast and mammalian cells (Peskett et al., 2018). While the soluble oligomers are recognized as major toxic agents due to their ability to aberrantly engage various cellular machineries (Bennett et al., 2005; Kim et al., 2016; Leitman et al., 2013; Miller et al., 2011; Park et al., 2013; Schaffar et al., 2004), the inclusions, though apparently less toxic (Arrasate and Finkbeiner, 2012; Arrasate et al., 2004; Slow et al., 2005), contribute to cytopathology by sequestering key cellular proteins and physically disrupting sub-cellular membrane structures (Bauerlein et al., 2017; Gruber et al., 2018; Kim et al., 2016; Park et al., 2013; Sánchez et al., 2003).

Mammalian cells, including neurons, employ two major pathways for the clearance of misfolded and aggregated proteins, the ubiquitin-proteasome system (UPS) and macroautophagy (hereafter autophagy). Stable aggregates are not directly accessible to the UPS for steric reasons (Bence et al., 2001; Guo et al., 2018; Holmberg et al., 2004; Venkatraman et al., 2004; Verhoef et al., 2002). Their clearance requires either chaperone-mediated disaggregation prior to proteolysis (Nillegoda et al., 2018; Saha et al., 2023; Tyedmers et al., 2010) or encapsulation by autophagosomes for lysosomal degradation (Hara et al., 2006; Iwata et al., 2005; Komatsu et al., 2006; Leeman et al., 2018; Mizushima, 2018; Ravikumar, 2002). The significance of the autophagy pathway is underscored by the finding that mutations in its core components are associated with neurodegenerative diseases, including mutations in p62/sequestosome (SQSTM1), optineurin (OPTN), and possibly ubiquilin 2 (UBQLN2), which is more closely linked to the UPS pathway (Deng et al., 2017; Hjerpe et al., 2016; Levine and Kroemer, 2019; Nixon, 2013; Stamatakou et al., 2020). Indeed, autophagy has been implicated in aggregate clearance in a range of neurodegenerative diseases (Boland et al., 2018; Hansen et al., 2018; Levine and Kroemer, 2019; Nixon, 2013; Rubinsztein et al., 2011), prominently including Huntington’s disease and other polyQ expansion disorders (Ashkenazi et al., 2017; Ravikumar, 2002; Sarkar et al., 2007). Autophagy of Htt inclusions involves recognition of ubiquitylated Htt (DiFiglia et al., 1997; Sap et al., 2019) by the autophagy adaptor protein p62, which forms a bridge between cargo and the protein MAP1LC3B (LC3B) anchored to the autophagosome membrane (Bjorkoy et al., 2006; Deng et al., 2017), and condenses the cargo through oligomerization (Wurzer et al., 2015). Downstream of the pathway, cargo-containing autophagosomes fuse with lysosomes for content degradation. Based on results from model systems, activation of autophagy by stimulating 5’ AMP-activated protein kinase (AMPK) with agents such as trehalose or inhibition of mTOR with rapamycin can ameliorate aggregate cytopathology in a number of model systems (Kim et al., 2011; Rusmini et al., 2019; Sarkar et al., 2007; Tanaka et al., 2004).

Exactly how aggregates are recognized and engulfed by autophagy has remained elusive. The process appears to be of limited efficiency and it has been suggested that polyQ aggregates are typically too large to be engulfed (Bauerlein et al., 2017). In this study, we employed cryo-correlative light and electron microscopy (cryo-CLEM) to visualize the engagement of Htt polyQ aggregates by autophagosomes *in situ*. We find that autophagy preferentially targets the amorphous polyQ phase, which interacts productively with p62. In contrast, the amyloid-like fibrils exclude p62 and evade phagophore engulfment. Solidification of the amorphous polyQ phase tends to trap the autophagic machinery. These results can explain the limited efficiency of autophagy in clearing solid polyQ aggregates.

## Results

### PolyQ aggregates varying in repeat length are differentially targeted by autophagy

To test the role of autophagy in the clearance of polyQ aggregates in neuronal cells, we expressed ecdysone-regulated Htt exon 1-GFP fusion proteins with repeats of 64Q and 150Q (Kim et al., 2016; Wang et al., 1999). After 48 hours of induction with muristerone A, protein aggregates of different sizes were visible by confocal fluorescence light microscopy (Figure S1A). Most 64Q appeared to be diffuse in the cell, with ∼10% of cells harboring small aggregates of varying brightness. In contrast, 150Q formed large GFP-intense globular structures that were previously observed to contain amyloid-like fibrils (Bauerlein et al., 2017). To monitor aggrephagy, we co-expressed the mammalian Atg8 homolog MAP1LC3B/LC3B as a marker, which is conjugated to phosphatidylethanolamine of the phagophore membrane during phagophore biogenesis. mCherry-LC3B co-localized with 64Q but not with the aggregates of 150Q (Figure 1A). Similarly, Lamp1-RFP, a lysosomal marker, also co-localized with 64Q but not with the 150Q aggregates (Figure 1B), suggesting a polyQ length-dependent targeting of polyQ aggregates by autophagy.

**Figure 1.**
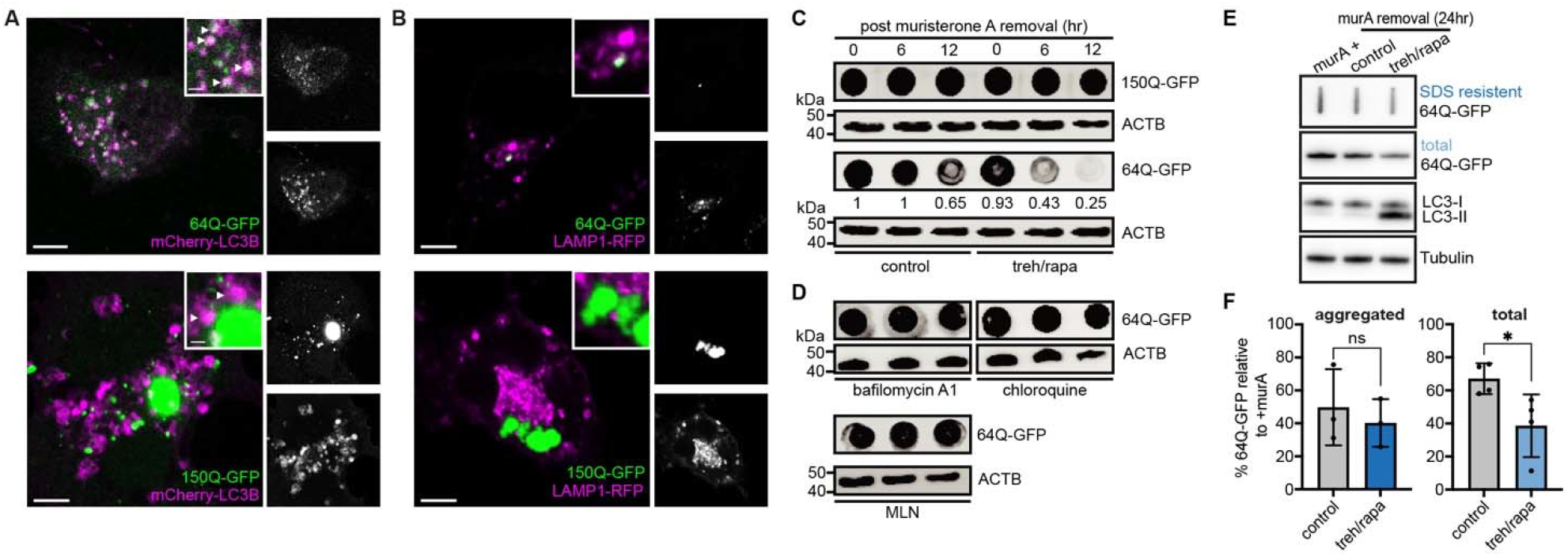
PolyQ aggregates with varying repeats are differentially targeted by autophagy. **(A and B**) Representative confocal microscopy images of 2-day muristerone A (1μM) induced expression of 64Q- and 150Q-GFP in Neuro2a, co-expressed with mCherry-LC3B (A) or Lamp1-mCherry (B). Arrowheads: polyQ-LC3B interactions. **(C and D)** Dot blots with the indicated antibodies for total 150Q- and 64Q-GFP lysates with 2-day muristerone A induction and removal, with a 1-day concurrent 150mM treh /200nM rapa treatment, or with autophagy inhibited by bafilomycin A (100nM) or chloroquine (100μM), or with Ub-activating enzyme E1 inhibited by MLN7243 (0.2uM) for 6 or 12 hours. ACTB as loading control. **(E and F)** Slot and immunoblots (E) with the indicated antibodies for aggregated or total 64Q-GFP with 2-day muristerone A induction and removal, followed by a 1-day 150mM treh /200nM rapa treatment, tubulin as loading control. Quantification with normalization to +muristerone A (100%) (F), error bars: s.d. n=3 for western blot, n=4 for filter-trap, *p<0.05.

To compare the autophagic clearance of polyQ aggregates with different polyQ lengths, we monitored the levels of total polyQ-GFP after 6 and 12 hours of muristerone A withdrawal by dot blot assay (Figure 1C). Consistent with the differential co-localization of 64Q and 150Q with the autophagic machinery (Figure 1A and 1B), only 64Q, but not the more fibril-prone 150Q, showed a visible reduction by immunoblotting after 12 hours of muristerone A withdrawal. To increase autophagic turnover, we treated the cells with a combination of trehalose and rapamycin (Sarkar et al., 2007), and monitored the efficacy of autophagy induction by analyzing lipidated LC3B-II levels (Figure S1B). A robust increase in LC3B-II levels was observed after 24 hours of treatment with 150 mM trehalose and 200 nM rapamycin (treh/rapa) without compromising cell viability (Figure S1B). Treh/rapa treatment enhanced clearance of 64Q aggregates but not 150Q aggregates (Figure 1C). In contrast, blocking lysosomal degradation with bafilomycin A or chloroquine (Mauthe et al., 2018) abolished 64Q degradation (Figure 1D), indicating that 64Q is indeed subject to autophagic degradation. Inhibition of the Ub-activating enzyme E1 by MLN7243 (Hyer et al., 2018) also abolished 64Q degradation (Figure 1D), suggesting that polyQ clearance is a ubiquitin (Ub)-dependent process.

The Htt polyQ model protein has been shown to undergo a liquid-to-solid phase transition that ultimately drives the aggregation process toward SDS-resistant amyloid fibrils, with the aggregates increasing in fluorescence intensity along this pathway (Peskett et al., 2018). To gain information about the structural state of the 64Q aggregates, we turned to *in situ* cryo-CLEM (Figure S1C-F) (Arnold et al., 2016; Goetz and Mahamid, 2020). In short, 64Q expressing cells were cultured on grids and vitrified by plunge freezing (Schorb et al., 2017). The GFP signal identified by the cryo-CLEM workflow allowed targeting of the aggregates during lamella preparation using focused ion beam (FIB) milling; tilt-series at correlated regions were subsequently collected by cryo-electron tomography (cryo-ET). In addition to the known fibrillar phase of 64Q (Bauerlein et al., 2017), this revealed the presence of a structurally amorphous non-fibrillar phase of 64Q. The amorphous 64Q phase corresponded to a dim GFP puncta in fluorescence microscopy, much smaller than the large aggregate signal predominantly seen in 150Q (Figure S1A and S1C). This amorphous phase contained many double membrane structures, some of which resembled potential phagophores and intermediates in autophagosome formation (Bieber et al., 2022; Li et al., 2023) (Figure S1F and S1G). Furthermore, biochemical analysis revealed a ∼50% decrease in the total pool of 64Q protein upon autophagy induction, although the amount of SDS-resistant aggregates was only slightly reduced (Figure 1E and 1F), indicating that autophagy preferentially targets non-fibrillar, SDS-soluble Htt aggregates.

Taken together, these data suggested that autophagy efficiently removes the SDS-soluble aggregates of 64Q, whereas autophagic clearance is limited for the SDS-resistant, fibrillar forms, predominantly seen with the longer 150Q repeats.

### Autophagy affects the different phases of 97Q aggregates

To genetically manipulate polyQ aggrephagy, we expressed 97Q-GFP (97Q) in HEK293 cells, aggregates of which have been shown by cryo-ET to form fibrils (Bauerlein et al., 2017). As judged by fluorescence light microscopy, 97Q and 150Q aggregates are similar in terms of brightness and prevalence. To monitor the size of the 97Q aggregates during autophagy, we co-labeled cells with either an antibody against LC3B or LysoTracker to stain lysosomes (Figure 2A and 2B). We quantified the cross-sectional areas around the equator of the aggregates as well as signals corresponding to LC3B or LysoTracker within a 1.5 μm distance from the aggregates (Schindelin et al., 2012). To downregulate autophagy, we performed siRNA knock down of LC3A/B/C (Weidberg et al., 2011), or CRISPR/Cas9 knock out of LC3B, as confirmed by immunoblotting (Figure S2A). Autophagy was again induced by treh/rapa treatment (24 hours), which resulted in a significant increase in the level of lipidated LC3B-II (Figure 2C and S2B) without impairment of cell viability as judged by live Annexin V staining (Figure S2C).

**Figure 2.**
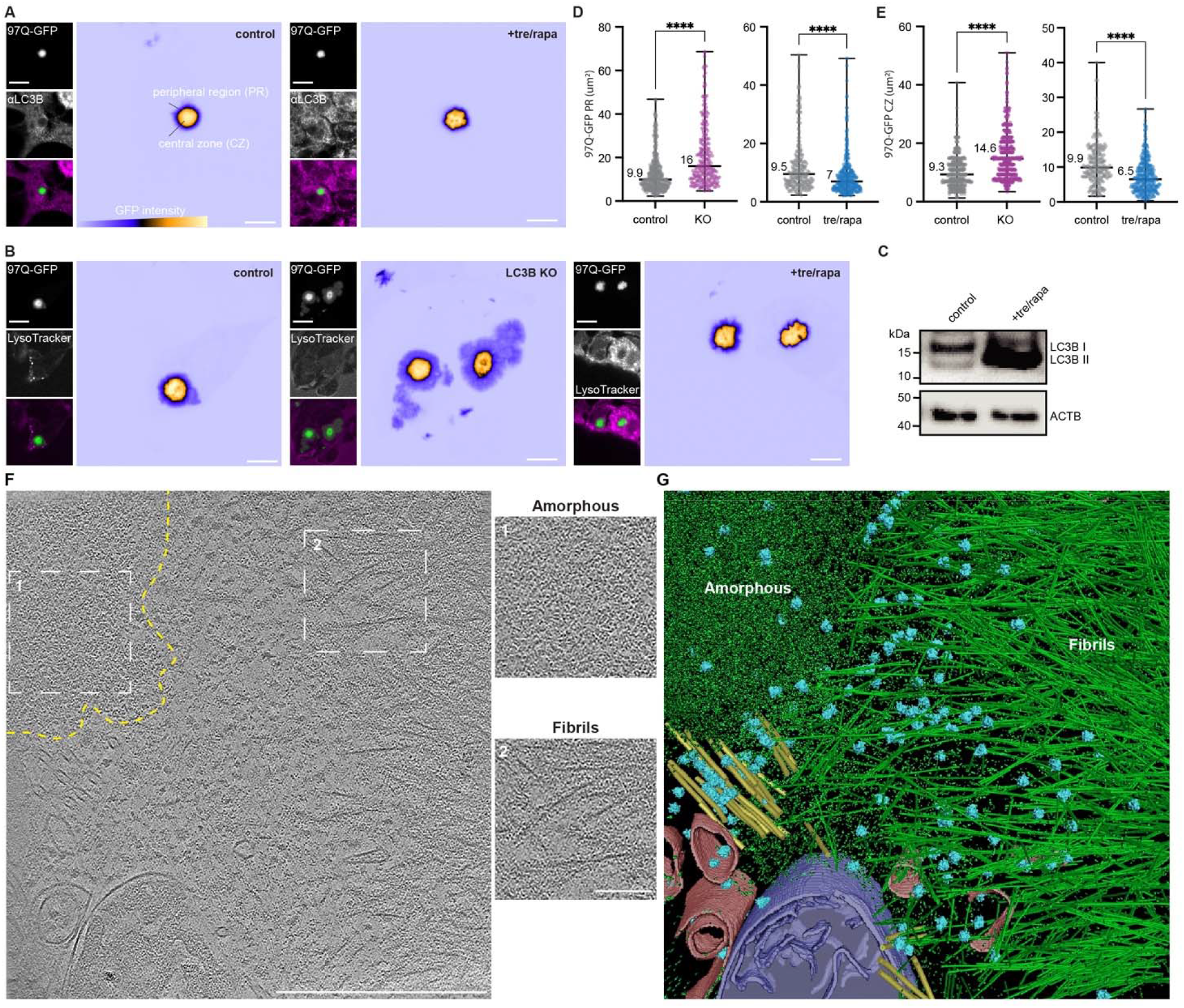
Autophagy affects the different phases of 97Q aggregates. **(A and B)** Representative confocal images of control, LC3B knock out (KO), or 1-day 150mM treh/200nM rapa treatment in HEK293, stained with LC3B antibody (A) or LysoTracker (B). **(C)** Immunoblot against LC3B for treh/rapa induced autophagy, ACTB as control. **(D)** Image quantification of 97Q-GFP peripheral region (PR) in control v. LC3B knock out (control n=395, KO n=245), v. treh/rapa (control n=228, +treh/rapa n=310) with medians displayed, intensity standardized with threshold 800-8000, ****p<0.0001. **(E)** Image quantification of 97Q-GFP central zone (CZ) cross-sectional area in control v. LC3B knock out (control n=409, KO n=551), v. treh/rapa (control n=226, treh/rapa n=308) with medians displayed, central zone intensity standardized with threshold >8000 (Fiji), ****p<0.0001. **(F and G)** Tomographic slice of 97Q-GFP at 32000x (F) and segmentation (G). ER related membranes (pink), mitochondria (purple), microtubules (yellow), ribosomes (blue), and polyQ (green). Inserts show enlarged amorphous (dotted line) and fibrillar regions. [CLEM workflow and lamella view: SI Figure 1A-C] Scale bars: 5μm in (A, B), 500nm in (F).

We observed that 97Q aggregates under constitutive expression consisted of two distinct zones: a central zone of high fluorescence intensity and a much dimmer peripheral region surrounding it (Figure 2A, 2B, and S2D). Upon LC3 knockdown or LC3B knockout, microscopic analysis consistently revealed an increase in the central zone (Figure 2E and S2E), coupled with a loss of LysoTracker signals around the polyQ aggregate (Figure S2F). In addition, the loss of LC3 resulted in an enlargement of the peripheral region around the central aggregate zone (Figure 2B, 2D, and S2E). The fluorescence intensity of the peripheral aggregate region under these conditions was only 10-25% of that of the central zone, indicating a substantially lower 97Q density. In contrast, induction of autophagy by treh/rapa reduced the size of the aggregates (Figure 2A and 2B), as manifested by a reduction in both the peripheral region and the central zone (Figure 2D and 2E) and an increase in the proximal LysoTracker and LC3B signals (Figure S2F and S2G), suggesting effects of induced autophagy during polyQ aggregate growth. To rule out effects of the GFP-tag on 97Q, we performed experiments with cells transfected with 97Q-myc, which resulted in similar observations of aggregate expansion upon knockdown of LC3A/B or LC3A/B/C (Figure S2H).

During aggregation multiple physical states of polyQ Htt are presumably in equilibrium, ranging from liquid-like condensates to solid, amorphous aggregates and end-stage amyloid-like fibrils (Bauerlein et al., 2017; Gruber et al., 2018; Peskett et al., 2018). We therefore examined the dynamics of 97Q in the central zone and peripheral region by fluorescence recovery after photobleaching (FRAP) (Figure S2I). FRAP analysis using a double normalization method showed that the mobile fraction of 97Q in the central zone was estimated at 0.34 ± 0.08 (mean ± SD) and remained unchanged upon induction of autophagy (0.31 ± 0.08), indicating that the majority of polyQ in the central zone is immobile regardless of the state of autophagy. In contrast, while the recovery of the peripheral region was similarly low after LC3B knockout (0.32 ± 0.1), the mobile fraction of 97Q more than doubled after treh/rapa induced autophagy (0.68 ± 0.2) or mCherry-LC3B overexpression (0.62 ± 0.3), indicating that activation of autophagy increases the mobile fraction within the peripheral region and counteracts its solidification.

To visualize the ultrastructural differences between the central zone and peripheral region of the polyQ aggregate, we again turned to *in situ* cryo-CLEM. Cryo-CLEM analysis of LC3B knockout cells expressing 97Q, showed that the high GFP intensity of the central zone corresponds to a polyQ fibrillar core, while the lower GFP signal of the peripheral region corresponds to an amorphous polyQ phase, both of which largely excluded ribosomes (Figure 2F, 2G, and S2J). In addition, amorphous polyQ appeared in two states: one contained some fibrillar structures within a confined amorphous density consistent with a more solid state and corresponding to a brighter GFP fluorescence (Figure S2J and SI1D-F); the other state lacked detectable fibrillar structures and corresponded to a weaker GFP signal, which nevertheless separated from the cytosol, as ribosomes were largely excluded from these areas (Figure 2F and 2G, S2J, and SI1). Ongoing liquid-to-solid phase transition within the amorphous polyQ phase may explain the low mobility of 97Q in the peripheral region observed by FRAP analysis (Figure S2I).

### Aggrephagy preferentially engulfs the amorphous polyQ phase

The results so far suggested that autophagy can only remove an early intermediate in the Htt aggregation pathway (Figure 1 and 2). We next used cryo-CLEM to study the interaction of the autophagy machinery with the aggregates (Figure 3, S3, SI 2-6). A schematic of the expected autophagic structures transitioning from cup-shaped phagophores to bilamellar autophagosomes, and fusion of the autophagosome with lysosomes to form autolysosomes for degradation is shown in Figure 3A. To target autophagic structures, cells expressing 97Q with mCherry-LC3B or after LysoTracker staining were used for the cryo-CLEM workflow. To ensure the identity of the structures in the lamella, the surviving thin lamellae were re-imaged in the cryo-confocal microscope after tomogram acquisition to post-correlate the fluorescent signals (Figure SI4C, SI4K and SI6C).

**Figure 3.**
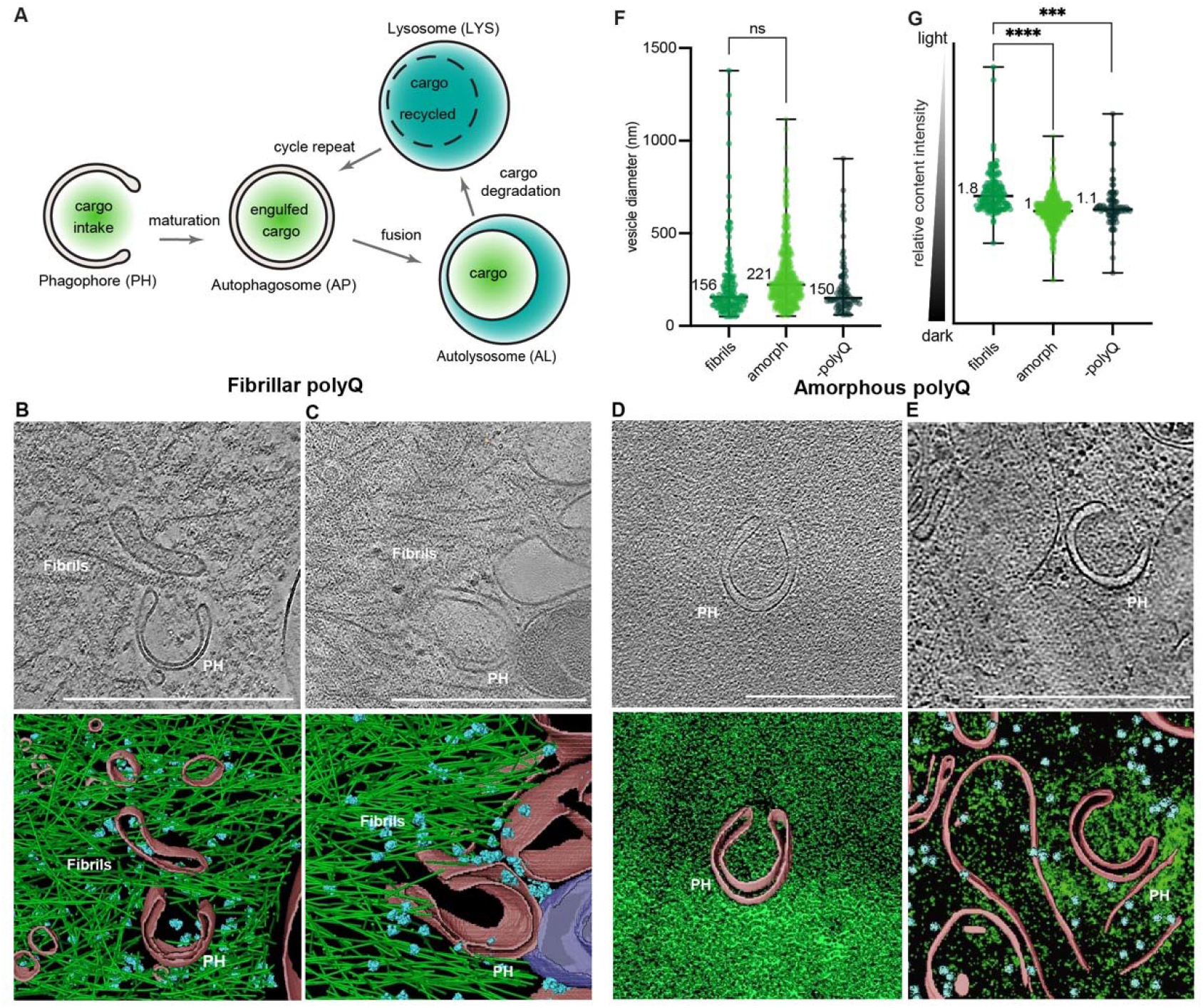
Phagophores preferentially interact with amorphous polyQ aggregates *in situ*. **(A)** Schematic of the different stages of the autophagic degradation pathway. **(B-E)** Tomographic slices acquired for 97Q-GFP with LysoTracker staining or mCherry-LC3B co-expression upon induced autophagy, for phagophores (PH) proximal to fibrils (B and C) or amorphous polyQ (D and E). Segmentation: PH and ER related membranes (pink), mitochondria (purple), ribosomes (blue), and polyQ (green). **(F and G)** Tomogram quantification: diameter (F) and content density (G) for phagophores to autophagosomes, proximal to only fibrils (n = 143), to amorphous polyQ (n = 584), or in cells without polyQ (n = 81) with median values, ****p<0.0001, ***p<0.001. [CLEM and lamella views: B (Figure SI 2), C (SI 3), D (SI 4I-K), and E (SI 4E-H)]. Scale bars: 500nm in (B-E).

Tomograms revealed different autophagic structures close to both fibrillar and amorphous polyQ phases, including phagophores (Figure 3B-E, S3E, and SI2-4, 6), autophagosomes, and autolysosomes with not-yet digested cargo content (Figure S3A-G, SI4-6). Often these structures appeared to be trapped and isolated in the observed polyQ phases. These trapped structures were morphologically aberrant compared to structures seen previously in cells not expressing polyQ proteins (Carter et al., 2020; Li et al., 2023). Small phagophores for example showed a strong cup-shaped bending and the intermembrane distance between the inner and outer bilayer of the phagophore membrane was often wider as seen in previous cases without polyQ expression. Notably, phagophores proximal to or trapped by fibrillar polyQ (Figure 3B and 3C), as well as autolysosomes and lysosomes (Figure S3A-D), were largely cargo-free, as judged by their electron transparent interior, indicating failed uptake of fibrillar aggregates. On the contrary, phagophores (Figure 3D and 3E) and their mature forms (Figure S3E-G) proximal to amorphous polyQ were filled with electron-dense cargo, resembling the surrounding amorphous density. These observations were supported by post-correlation analysis, demonstrating that the lamella regions with the autophagic structures were indeed positive for both 97Q and mCherry-LC3B (Figure S3E, S3F, SI4D, and SI6D). Strikingly, trapped phagophores were also observed in a dense amorphous polyQ phase devoid of other cellular components, suggesting that a potential solidification of the polyQ phase may block phagophore maturation (Figure 3D, SI4H). To further test this possibility, we co-transfected cells with 97Q-myc and the autophagic flux reporter GFP-LC3B-RFP (Kaizuka et al., 2016). The GFP-LC3-RFP reporter is cleaved in the linker between LC3 and RFP by the activating enzyme Atg4, resulting in equimolar amounts of GFP-LC3 and RFP. While GFP-LC3 is lipidated and subsequently degraded during autophagy, RFP serves as an internal expression control, allowing the determination of autophagic flux based on the GFP/RFP signal ratio. Strikingly, this analysis revealed that the reporter signals around the large aggregate core were green-shifted (Figure S3H) and completely immobile for an extended period of time during time-lapse microscopy (Figure S3I), indicating a block of autophagic turnover. Several examples of ER-phagy were also observed proximal to the aggregates (Figure S3E and S3G), containing ER cargo but again lacking fibrillar polyQ content.

Using ∼300 tomograms, we categorized vesicular membrane structures of the autophagic pathway (corresponding to phagophores, autophagosomes, autolysosomes, and lysosomes) (Figure 3F, 3G S3J, and S3K) as either proximal to fibrils or to amorphous polyQ, or in cells without polyQ overexpression. These vesicular membrane structures were then analyzed for their dimensions and cargo density. No significant size difference was observed for vesicular membrane structures in the three categories, as they covered a wide range of vesicle diameters (Figure 3F). However, the interior of vesicular membrane structures proximal to fibrils was significantly less dense, as quantified with respect to the density of the entire tomogram (Figure 3G), suggesting absence of cargo. The downstream vesicles in the vicinity of fibrils also showed a similar lack of cargo engulfment (Figure S3K); they were smaller than those proximal to amorphous polyQ, possibly due to fibril-induced deformation (Figure S3J). In conclusion, *in situ* cryo-CLEM indicates that autophagic structures preferentially interact with and engulf the amorphous phase of polyQ aggregates and not the fibrillar form.

### Autophagic targeting of polyQ is a p62-mediated process

Recently, the CCT2 subunit of the chaperonin TRiC/CCT was reported to serve as a Ub-independent selective autophagy receptor for the removal of solid polyQ aggregates (Ma et al., 2022). To identify the autophagy receptor for polyQ in an unbiased manner, we established a protocol to isolate polyQ-containing autophagosomes and analyze their contents by label-free, quantitative mass spectrometry (Cox et al., 2014). Vesicles were isolated from LC3B knockout cells co-expressing 97Q and mCherry-LC3B after addition of chloroquine and subjected to fluorescence sorting (Gao et al., 2010) (Figure 4A). Fluorescence sorting revealed that 10% of the total mCherry-LC3B vesicle pool in double-transfected cells were positive for both mCherry-LC3B and 97Q, indicating that the cells have a modest basal autophagic activity for polyQ. Induction of autophagy by treh/rapa treatment increased this percentage to 25%, indicating enhanced autophagic uptake of 97Q (Figure 4B and 4C). Next, the sorted non-fluorescent (negative), mCherry-positive (mCherry+), and the mCherry/GFP double-positive vesicles (mCherry+/GFP+) were examined in dot blot experiments to confirm the presence of LC3B and polyQ by immunoblotting (Figure 4D). The sorted vesicles from both cell systems (HEK293 and Neuro2a) were then analyzed by mass spectrometry (Figure 4E, 4F, S4A, and S4B).

**Figure 4.**
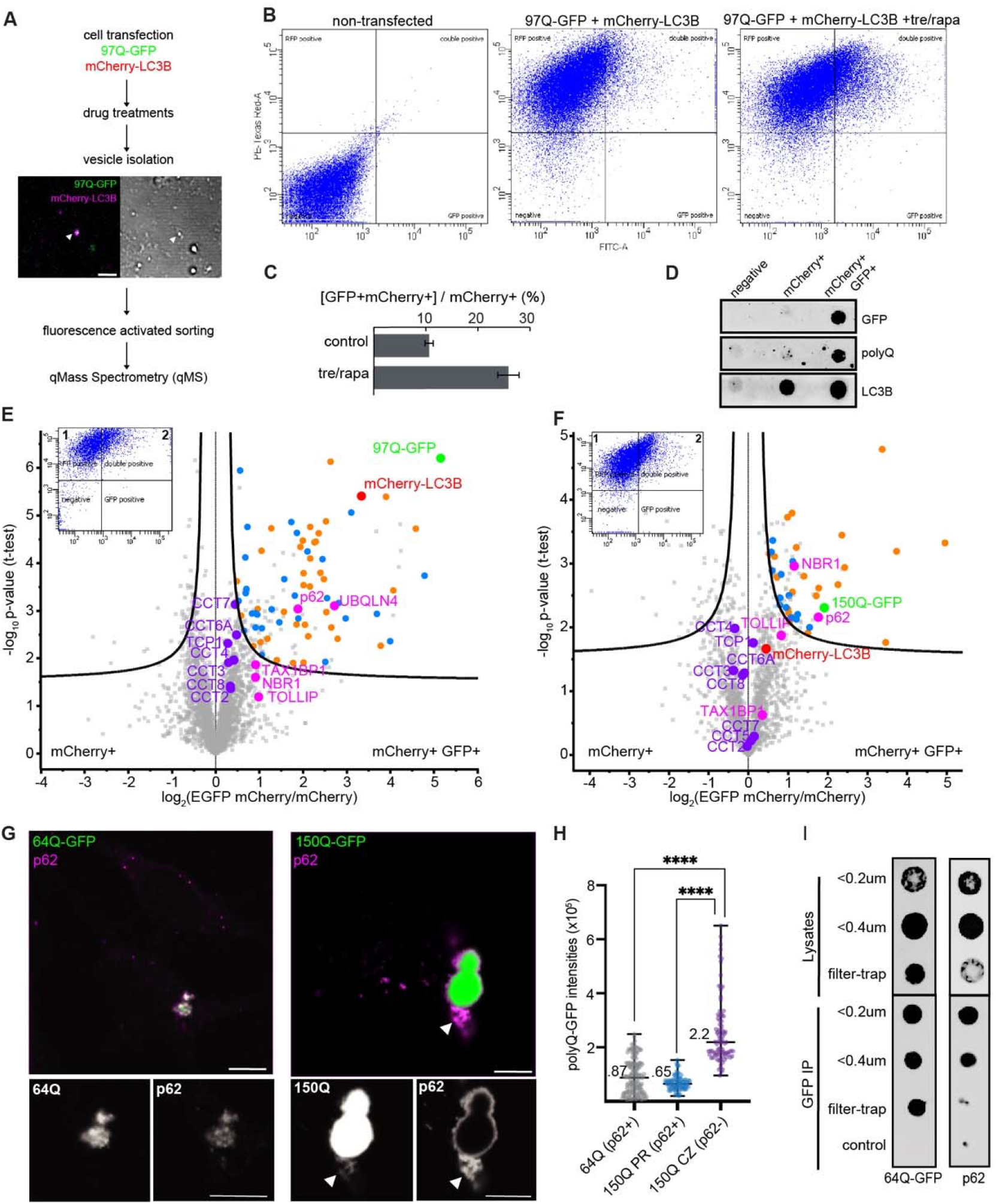
The autophagic intake of the polyQ is a p62 mediated process. **(A)** Experiment workflow: mCherry-LC3B and 97Q-GFP positive vesicles (arrowheads) extracted from co-transfected LC3B knock out. **(B-D)** Fluorescence vesicle sorting (∼10,000 events) sorted by RFP (Y axis) and GFP (X axis), puncta represent individual events (B), quantified as ratio of mCherry+GFP+ / mCherry+, error bars: n = 3, s.e.m (C). Sorted vesicles validated with the indicated antibodies in dot blots (D). **(E and F)** Label-free quantitative mass spectrometry analyses of the sorted vesicles plotted with mCherry+ control v. mCherry+ GFP+. Curves (FDR < 0.01) for HEK293 + treh/rapa + chloroquine (E) and Neuro2a + treh/rapa + chloroquine (F). Inserts: vesicles sorting for mass spectrometry with mCherry+ (quadrant 1) and mCherry+ GFP+ (quadrant 2). Receptors for Ub-substrates (pink) and TRiC subunits (purple) are highlighted; proteasome and protein quality control regulators (orange), and RNA processing factors (blue) are colored. Chloroquine was included to inhibit degradation of autophagosomal contents for mass spectrometry. **(G)** Representative confocal images of 64Q- and 150Q-GFP stained with antibody against p62. Arrowheads: polyQ-p62 co-localization. **(H)** Image quantification of 64Q- (n=148) and 150Q-GFP (n=87) for GFP intensities with p62 co-localization, with medians displayed, CZ: central zone, PR: peripheral region ****p<0.0001. **(I)** anti-GFP pull down using 64Q-GFP lysates partitioned through syringe filter (0.2-0.4 μm), for the detection with the indicated antibodies in each fraction in dot blots. Scale bars: 2 μm in (A), 5 μm in (G).

The mCherry+/GFP+ vesicles from HEK293 or Neuro2a cells were strongly enriched for polyQ-GFP (35 or 5-10-fold, respectively) in comparison to the mCherry+ control vesicles (Figure 4E, 4F, S4A, and S4B). From both cell types, quantitative mass spectrometry analyses revealed a significant enrichment of the Ub-dependent autophagy receptor p62 with polyQ, especially upon chloroquine treatment (FDR < 0.01) (Figure 4E and 4F). Numerous proteasome subunits, regulators of protein quality control, and proteins with low complexity sequences (e.g. FUS, TARDBP), previously found to interact with polyQ (Kato et al., 2012; Kim et al., 2016), were also enriched (Figure 4E, 4F, S4A). However, there was no enrichment of the proposed aggrephagy receptor CCT2 (Ma et al., 2022) compared to subunits of the TRiC complex or the entire quantified autophagosome proteome.

Next, we validated the roles of Ub and p62 in polyQ autophagy (Figure 4G-I, S4C-G). The requirement for ubiquitylation in polyQ degradation was confirmed using treatment with the E1 inhibitor MLN7243 (Figure 1C), which resulted in a size increase of the 97Q aggregates compared to control cells (Figure S4C). While Ub antibody stained both the peripheral region and the central zone of the polyQ aggregate upon immunofluorescence analysis (Figure S4D and S4E), p62 was only detected in the dim 64Q aggregates and the peripheral region around the 150Q or 97Q aggregates (Figure 4G and S4F), suggesting a specific interaction of p62 with the amorphous polyQ phase. To exclude an experimental bias due to antibody staining, we co-expressed a fluorescent tagged version of p62, p62-RFP, confirming these observations (Figure S4G). The 64Q aggregates that interacted with p62 were indeed similar in GFP intensity to the 150Q peripheral region (Figure 4H), defining them as amorphous and thus amenable to aggrephagy through p62 binding. To further test the targeting of p62 to the amorphous polyQ, we isolated and partitioned cellular 64Q aggregates from cell lysates using filtration and subsequent enrichment by GFP pull-down. These experiments confirmed that p62 preferentially binds to the smaller aggregates but was not detected in the filter-trapped fraction (Figure 4I). These results demonstrate that p62 interacts with the amorphous non-fibrillar polyQ.

### Autophagosomes preferentially contain amorphous 97Q

Like all amyloids, polyQ fibrils consist of highly stable cross-β structure (Hoop et al., 2016; Perutz et al., 1994; Scherzinger et al., 1997). To provide further evidence that autophagosomes indeed preferentially engulf amorphous polyQ, we employed cryo-CLEM to examine the structural details of 97Q within autophagic vesicles. To this end, we plunge froze the fluorescence-sorted mCherry/GFP double-positive vesicles (mCherry+/GFP+) isolated from 97Q/ mCherry-LC3B expressing HEK293 cells after induction of autophagy (Gao et al., 2010). The vitrified vesicles were examined in a cryo-confocal fluorescence microscope to locate GFP and mCherry double-positive puncta; the z-stacks containing the fluorescence signals were then correlated with the cryo-TEM overviews to locate the vesicle for tilt-series acquisition (Figure 5A). This analysis revealed different autophagic vesicles, ranging from bilamellar autophagosomes (Figure 5A, 5B, and S5A) to unilamellar autolysosomes (Figure 5C and S5B). From the 200 autophagic vesicles examined (Figure 5D), only one mCherry+/GFP+ autophagosome contained some fibrillar material of uncertain origin, mixed with amorphous content (Figure 5B), while the rest of the autophagosomes and autolysosomes contained only amorphous density (examples in Figure 5A, 5C, S5A, and S5B).

**Figure 5.**
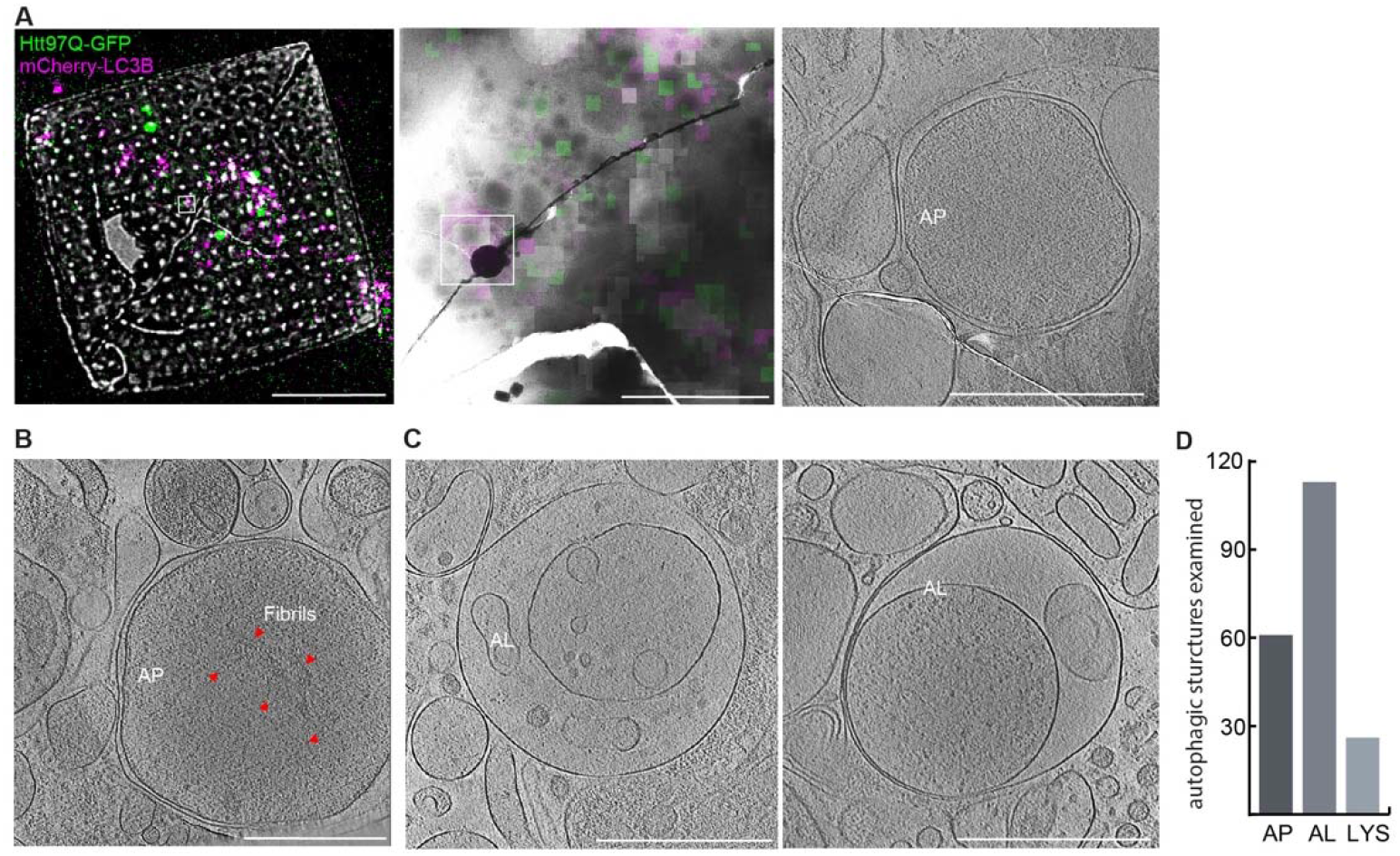
Autophagy preferentially intakes the amorphous phased 97Q. **(A)** Left: A grid square of vitrified vesicles from mCherry-LC3B and 97Q-GFP co-expressed LC3B knock out cells, imaged in the cryo-confocal with the tomogram site boxed. Middle: TEM image (5600x) with the corresponding tomogram site boxed overlayed with fluorescence. Right: tomographic slice (34000x) with the GFP positive autophagosome (AP) filled with amorphous content. **(B)** A GFP positive autophagosome that contains fibril-like structures. **(C)** A GFP positive autolysosomes (AL) with amorphous content. **(D)** Numbers of autophagic structures observed by cryo-ET upon isolation. Scale bars: 10μm (A left), 2μm (A middle) 500nm in (A-C tomographic slices).

Taken together, our results demonstrate that aggrephagy mediated by p62 preferentially targets amorphous rather than fibrillar polyQ aggregates.

## Discussion

The biogenesis of an autophagosome is a slow process that takes about 10 minutes (Geng et al., 2008). During this time autophagic cargo must be concentrated and segregated into a distinct entity that can be engulfed by the autophagosomal membrane. Multiple interactions between autophagy receptors and Atg8/LC3/GABARAP family members are needed to establish a high avidity interaction platform between the cargo and the phagophore, ensuring autophagosome biogenesis at the site of cargo recognition (Kirkin and Rogov, 2019). The physical properties of the cargo/receptor complex therefore should determine the efficiency of the autophagy process. In this study, we define the role of autophagy in polyQ degradation by visualizing polyQ aggregates within autophagosomes using cryo-CLEM. This structural analysis reveals that autophagy preferentially targets an amorphous polyQ phase rather than the fibrillar state (Figures 5A-B), proposing a model in which autophagy targets early stages of the polyQ aggregation pathway (Figure 6).

**Figure 6.**
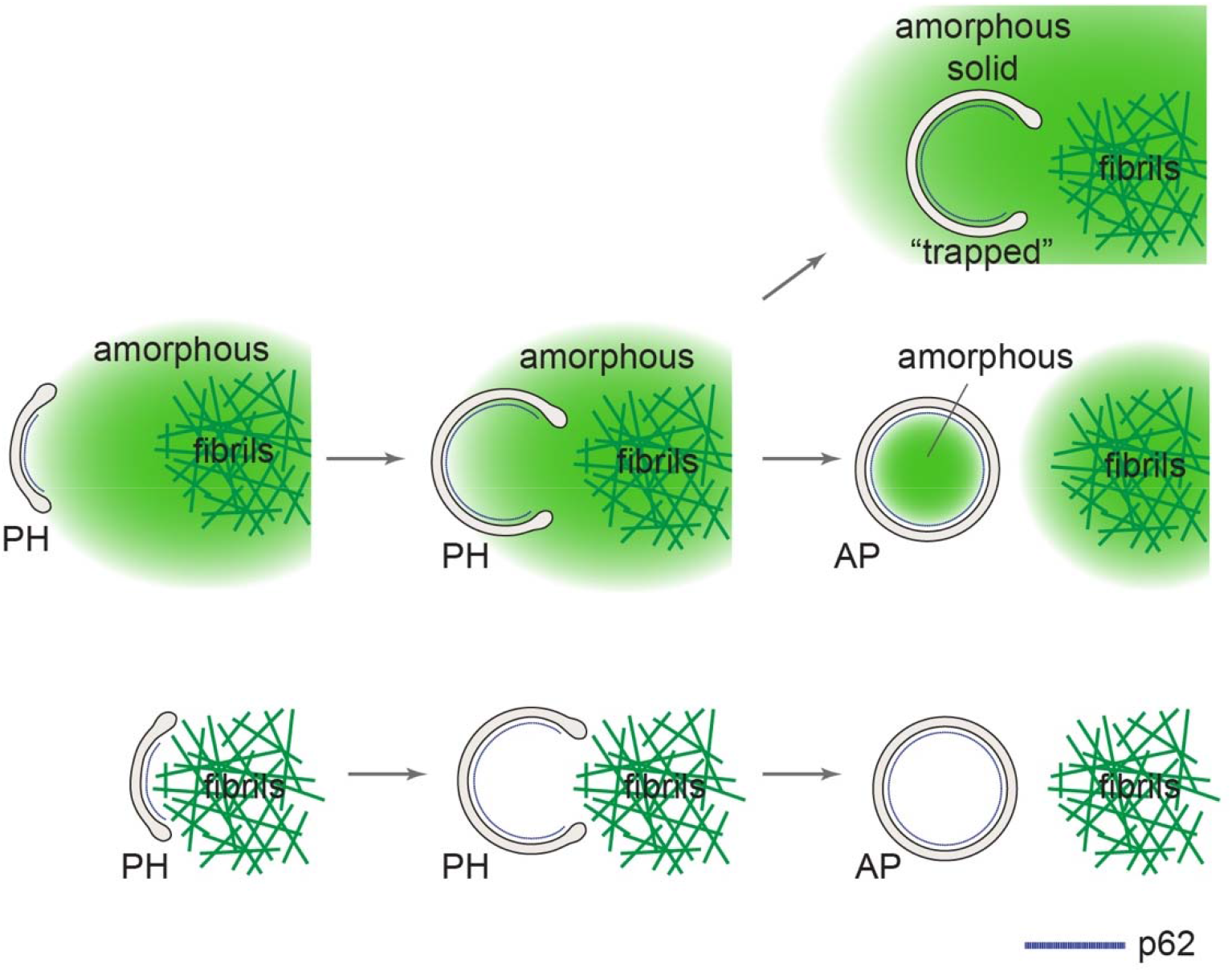
Model of p62 mediated autophagy of polyQ aggregates. Autophagy preferentially interacts and engulfs the amorphous polyQ phase mediated by the autophagy receptor p62. Amyloid fibrils show no obvious engulfment by autophagy and autophagic structures in close proximity are often observed to be cargo free. Both the fibrillar and the amorphous polyQ aggregates can trap phagophores leading to isolation and a block in phagophore maturation.

Recent reports have shown that various autophagic cargoes together with their corresponding autophagy receptors undergo co-condensate formation in the cytosol prior to autophagic engulfment (Wilfling et al., 2020; Yamasaki et al., 2020; Zaffagnini et al., 2018; Zhang et al., 2018), highlighting a role of liquid-liquid phase separation in cargo recruitment. Furthermore, evidence has been presented that surface adhesion of liquid droplets (also known as wetting) is sufficient for liquid-like cargo/receptor condensate engulfment by phagophore membranes (Agudo-Canalejo et al., 2021). PolyQ aggregates initially exist in a liquid-like state but undergo a liquid-to-solid phase transition that shifts the equilibrium toward amyloid fibrils, with longer polyQ repeats undergoing more rapid transition (Peskett et al., 2018). Whether the amorphous polyQ aggregates that we identified as being degraded by autophagy is in a liquid-like or gel-like state remains to be determined. However, quantitative mass spectrometry experiments identified p62 as the major autophagy receptor in the polyQ-containing autophagosomes (Figure 4E and 4F). p62 is a ubiquitin-dependent autophagy receptor that forms phase-separated condensates in the presence of polyubiquitylated cargo molecules (Jakobi et al., 2020; Sun et al., 2018; Zaffagnini et al., 2018). Fluorescence imaging showed that p62 localized exclusively to less concentrated polyQ aggregates, corresponding to the amorphous polyQ phase (Figure 4G). It is likely that targeting of p62 to the amorphous polyQ phase coverts the polyQ aggregate to a more liquid-like state. Consistent with this hypothesis, FRAP analysis revealed a larger mobile fraction of 97Q within the amorphous polyQ phase when autophagy was activated by treh/rapa or during LC3B overexpression, compared to LC3B knock out cells (Figure S2I).

In contrast, Ub was found in both the amorphous and fibrillar polyQ (Figure S4D and S4E), raising the question how p62 targets specifically the amorphous polyQ phase. Recently, it has been shown that the aggregate interface is primed for p62 binding by the protein NEMO, which amplifies linear ubiquitination by the E3 ligase HOIP and generates a mobile, phase-separated aggregate surface (Goel et al., 2023; Nikolas Furthmann and Gisa Ellrichmann, 2023; van Well et al., 2019). Our mass spectrometry analysis of isolated autophagosomes did not identify NEMO, suggesting that its interaction with the aggregate surface is transient.

Among the various ubiquitin-dependent autophagy receptors that function in selective autophagy, we find p62 to be of particular importance in polyQ aggrephagy, presumably based on its ability to phase-separate in the presence of ubiquitylated polyQ and link it to the phagophore-conjugated LC3BII (Bjorkoy et al., 2006; Wilfling et al., 2020; Yamasaki et al., 2020; Zaffagnini et al., 2018; Zhang et al., 2018). It is plausible that other receptors with domains important for oligomerization, such as TAX1BP1, NDP52, and NBR1 (Kirkin et al., 2009a; Lu et al., 2017; Nthiga et al., 2021), also prefer droplet-like cargo assemblies with a “soft” surface in mediating selective autophagy. It has been shown that NBR1 enhances p62-ubiquitin condensate formation and TAX1BP1 facilitates recruitment of the autophagy scaffold protein FIP200, suggesting that multiple factors may modulate the mobility of the amorphous polyQ phase (Jakobi et al., 2020; Kirkin et al., 2009b; Sanchez-Martin et al., 2020; Turco et al., 2021; Zaffagnini et al., 2018). A liquid-like state of the amorphous polyQ phase would provide a flexible surface for phagophore formation establishing high avidity, possibly resulting in piecemeal polyQ uptake (Figure 6), as seen previously for p62 droplets (Agudo-Canalejo et al., 2021). Such a mechanism would give rise to autophagosomes variable in size (Figure 3F) but generally much smaller than a full-sized aggregate deposit.

Contrary to a recent report (Ma et al., 2022), our study does not support the proposed role of the chaperonin subunit CCT2 as a selective receptor of polyQ fibril aggrephagy. Specifically, we failed to obtain evidence by mass spectrometry for a specific enrichment of CCT2 in polyQ-containing autophagosomes. Moreover, our cryo-CLEM analysis provides no evidence that amyloid-like fibrils are the main target during aggrephagy. Only in a single case did we observe fibril-like structures, of uncertain origin, surrounded by amorphous density within a polyQ positive autophagosome (Figure 5B). Nevertheless, our data raise the possibility that the fragmentation of fibrils by chaperones and their partitioning into an amorphous phase may contribute to aggregate clearance via autophagy.

Our experiments also showed that aggrephagy is not always productive and often leads to the trapping of phagophore membranes within the amorphous polyQ phase (Figure S3H and S3I). The trapped double membrane structures do not show the characteristic features of phagophores and are often isolated from other organelles (Figure 3B-E) (Bieber et al., 2022; Li et al., 2023). We speculate that the trapping of phagophores occurs due to a shift in polyQ material properties from liquid-like to solid during autophagosome biogenesis, incorporating the phagophore within the polyQ aggregate. Although the polyQ inclusions are apparently less toxic (Arrasate et al., 2004; Slow et al., 2005), they contribute to cytopathology by sequestering key cellular proteins and physically affecting the integrity of membranes (Bauerlein et al., 2017; Kim et al., 2016). The ensnaring of autophagic structures within both the amorphous and fibrillar phases of the polyQ aggregates highlights the detrimental effect of aggregate formation on the proteostasis network. Impaired autophagy may contribute to the severe pathology of longer polyQ repeats that convert to fibrils more rapidly (DiFiglia et al., 1997; Gusella and MacDonald, 2006). Thus the success of autophagy-based therapeutic interventions in polyQ diseases may depend on early detection prior to fibrillation or on ways to bypass a requirement for the Ub-p62 interaction (Li et al., 2019).

## Acknowledgements

We acknowledge Martin Spitaler and Markus Oster at MPI Imaging facility, Dr. Barbara Steigenberger and Nicole Krombholz at MPI Mass spectrometry facility for technical supports. We thank Dr. Dieter Edbauer for the p62-RFP construct. We acknowledge Dr. Jürgen Plitzko, Victoria Trinkhaus, Hou Zhen, Florian Beck, Dustin Morado, Guenter Pfeifer, Inga Wolf for technical support. We acknowledge Brenda A. Schulman for discussions and advice on the study. D.Y.Z acknowledges postdoctoral fellowship funded by the Alexander von Humbolt Foundation. This study was supported by the Max Planck Gesellschaft and funded in part by Aligning Science Across Parkinson’s ASAP-000282 (Brenda A. Schulman, F. Ulrich Hartl and Wolfgang Baumeister) through the Michael J. Fox Foundation for Parkinson’s Research (MJFF).

## Authors Contributions

D.Y.Z., F.W., W.B., F.J.B, and F.U.H designed the research project, W.B. provided instruments, D.Y.Z. and I.S. performed experiments with support from everyone. D.Y.Z, F.W., and F.U.H wrote the manuscript with input from the other authors.

## STAR Methods

### Key resources table

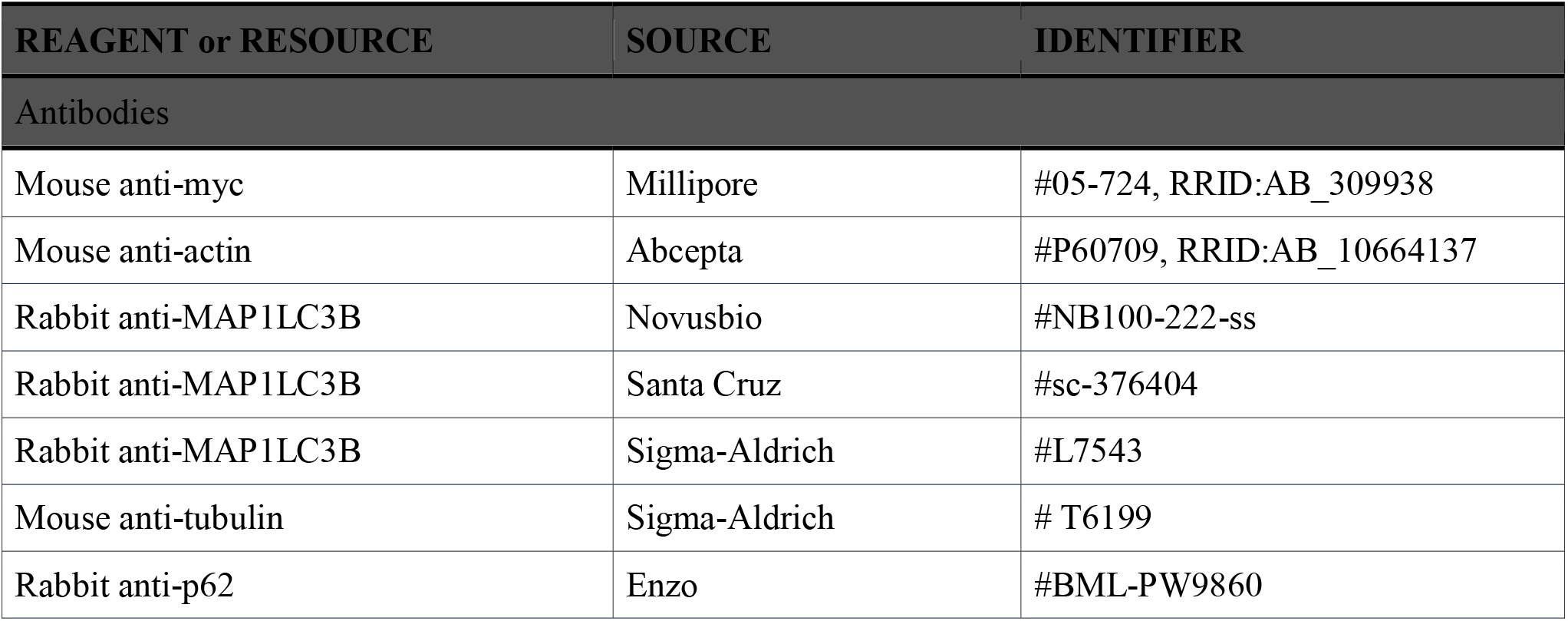
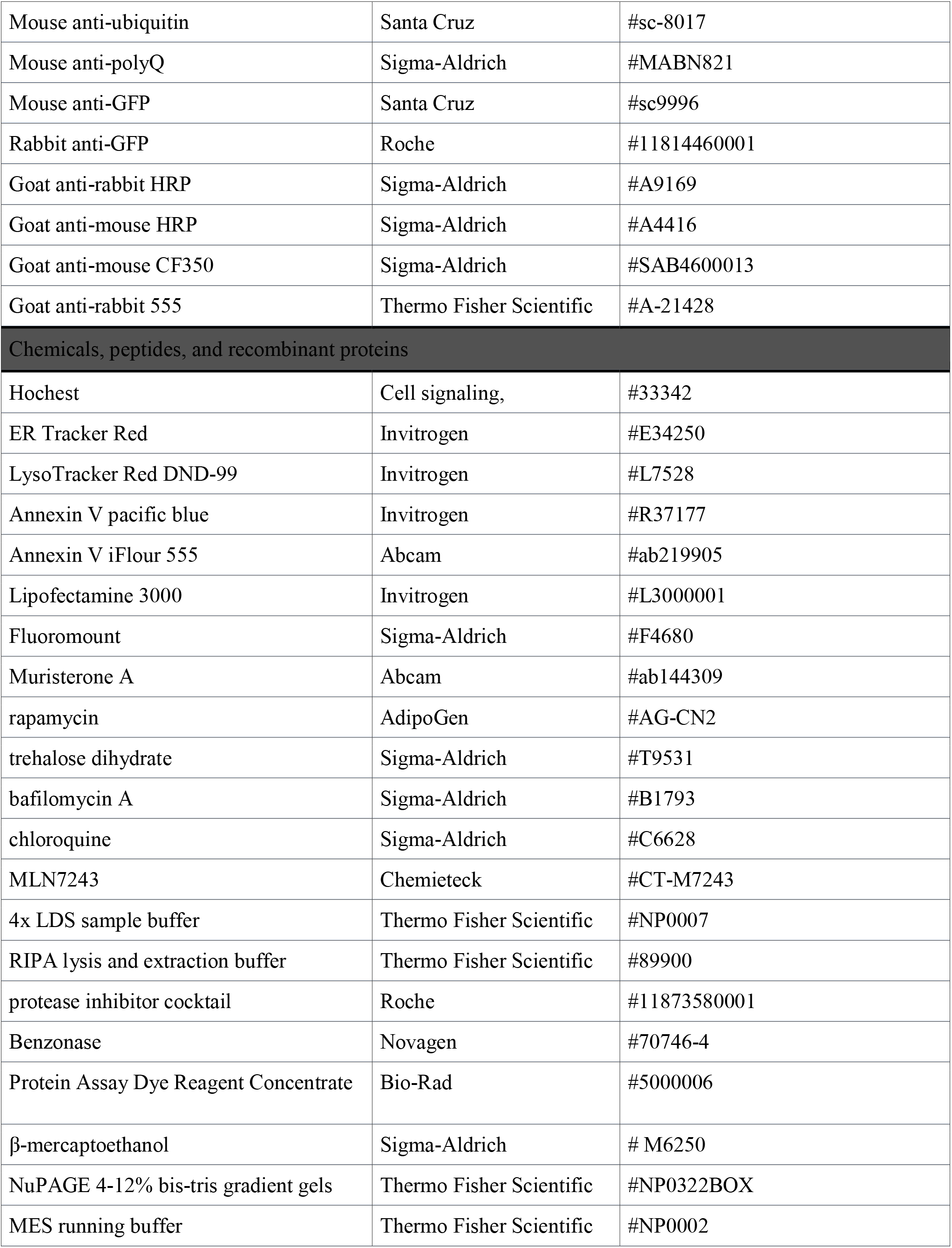
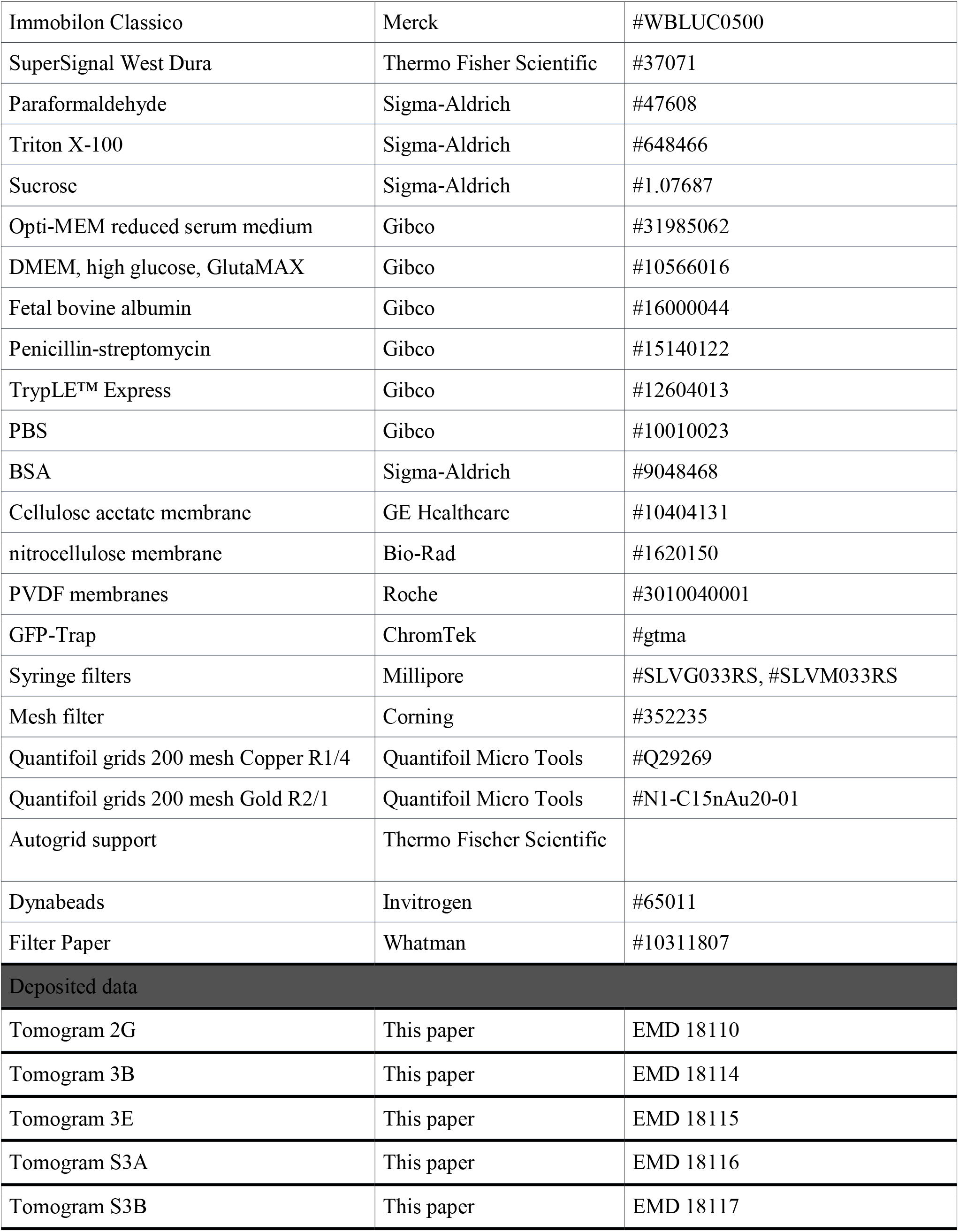
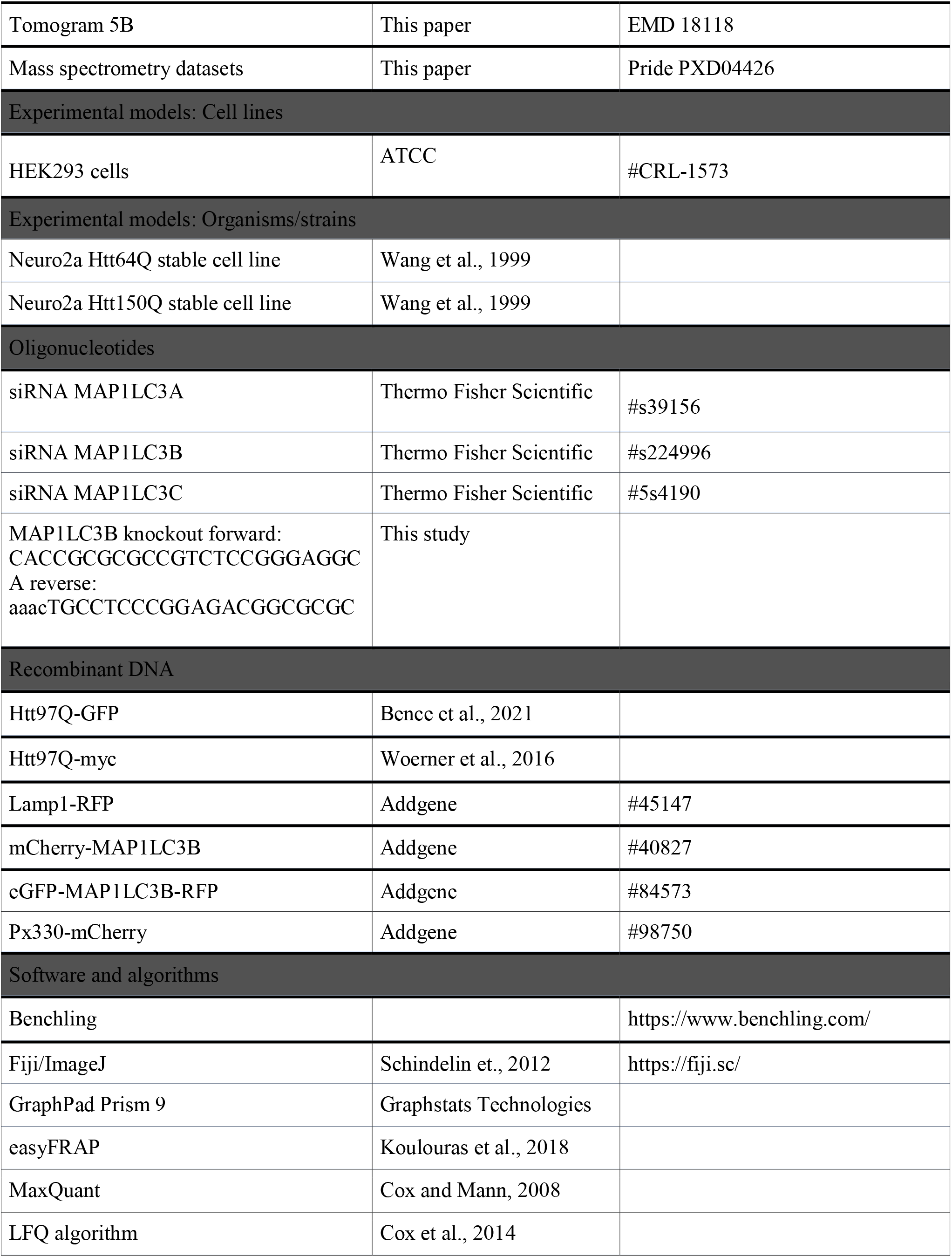
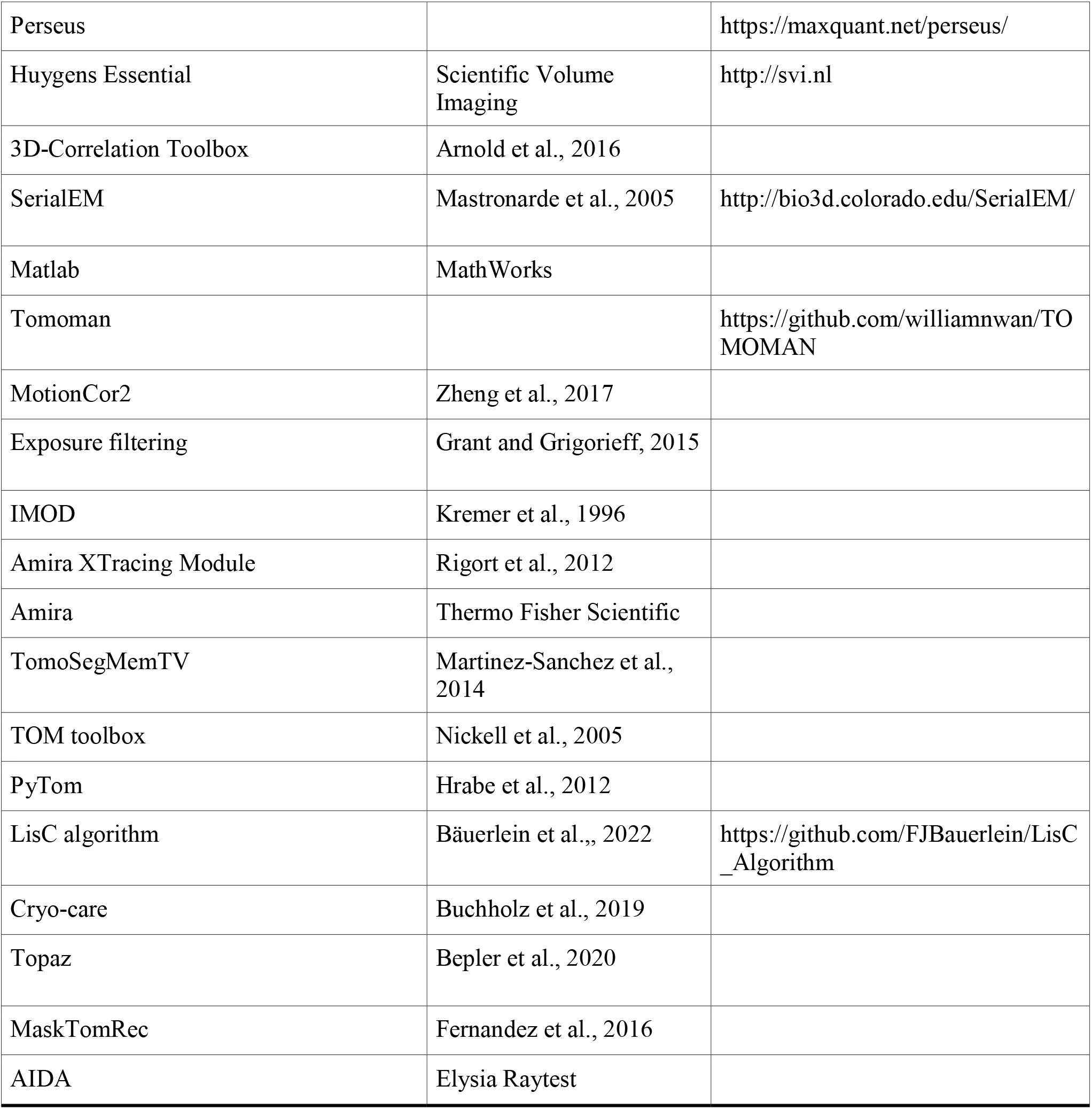

### Experimental Model and Subject Details

#### Cell lines, Plasmids, siRNAs, CRISPR/Cas9 knock out, and Chemicals

Cells were grown with DMEM + Glutamax (Gibco) with 10% FBS (Gibco) and 100 units/ml Penicillin-Streptomycin (Gibco) in incubator at 37°C with 5% CO2; no unusual Hoechst staining (Cell signaling) observed for mycoplasma contamination. For passages, cells were washed in PBS (Gibco), trypsinized with TrypLE™ Express (Gibco). Neuro2a containing stable expression of the inducible 64Q- or 150Q-GFP were described before (Kim et al., 2016). Early passages in which the 64Q had a higher propensity to form small-dim-amorphous aggregate over the bright fibrillar form (∼10:1) was used. For the induction of the transgene expressing polyQ, muristerone A (Abcam) was applied at 1 μM for 2 days. The following chemicals were applied to cell culture: rapamycin (AdipoGen), trehalose dihydrate (Sigma-Aldrich), bafilomycin A (Sigma), chloroquine (Sigma), MLN7243 (Chemieteck).

For transient expression, HEK293 were transfected for 48 hours using Lipofectamine 3000 (Invitrogen) in Opti-MEM (Gibco) as per manufacturer’s protocol, media was refreshed after 24 hours. The plasmids expressing GFP and myc tagged Htt97Q exon 1 were described previously (Bence et al., 2001; Woerner et al., 2016). Lamp1-RFP, mCherry-MAP1LC3B, eGFP-MAP1LC3B-RFP, and MAP1LC3B-3xflag were obtained from Addgene.

siRNA targeting human MAP1LC3A, B, and C and control were obtained from Thermo Fisher Scientific; 2-day transient knock-down was performed using Lipofectamine. CRISPR/Cas9 knock out of MAP1LC3B in HEK293 was generated with the px330-mCherry construct with the primer guides designed in Benchling (forward: CACCGCGCGCCGTCTCCGGGAGGCA and reverse: aaacTGCCTCCCGGAGACGGCGCGC). 2-day post transfection, individual positive clones were isolated through fluorescence-activated sorting with a 100 µm nozzle on BD FACSAria III using FACSDiva 6.1.3 software (MPI imaging facility, Martinsried, Germany). The knock down and knock out efficacies were assayed by immunoblotting.

## Method Details

### Immunoblotting and filter trap slot blots

The following antibodies were used for immunoblotting: mouse anti-GFP (Santa Cruz), rabbit anti-GFP (Roche), anti-MAP1LC3B (Sigma, Santa Cruz), anti-beta-actin (Abcepta), Anti-Tubulin (Sigma), Goat anti-rabbit HRP (Sigma), Goat anti-mouse HRP (Sigma).

Cells from a well of 12-well plate were lysed on ice in 80 µL RIPA buffer (Thermo) supplemented with protease inhibitor cocktail (Roche) and Benzonase (Novagen) with intermittent vortexing. Protein concentration in total cell lysates was determined using Protein Assay Dye Reagent Concentrate (Bio-Rad), normalized before denaturing in 4x LDS sample buffer (Thermo) containing 2.5% β-mercaptoethanol (Sigma) and boiling at 95 °C for 5 min. Proteins were separated on NuPAGE 4-12% bis-tris gels (Thermo) with MES running buffer (Thermo). Afterwards, proteins were transferred to PVDF membranes (Roche) in tris-glycine buffer using semi-dry transfer. Membranes were washed in TBS-T and blocked in 5% low-fat dry milk dissolved in TBS-T for 1 hour at room temperature. Blots were incubated with primary antibodies (1:500-1:1000) overnight at 4 °C, washed 3 times with TBS-T and probed with HRP-conjugated secondary antibodies (1:10000) for 1-2 hours at room temperature. Chemiluminescence was developed using HRP substrate Immobilon Classico (Merck) or SuperSignal West Dura (Thermo), detected on an ImageQuant800 (Amersham) imager with control software v1.2.0. Intensity of protein bands was quantified using AIDA image software v4.27.039 (Elysia Raytest).

Filter-trap assay for the detection of aggregated HttQ64-GFP was performed as before (Saha et al., 2023). Lysates were prepared as described in immunoblotting. 10 or 20μg total protein was diluted in 100µL or 200μL RIPA. Cellulose acetate membrane (0.2μm pore, GE Healthcare) was equilibrated in 0.1% SDS/H_2_O and fixed to the filter trap device (PR648 Slot Blot Blotting Manifold, Hoefer). Samples were loaded under vacuum. Slots were then washed with 200μL 0.1% SDS/H_2_O 3 times followed by standard immunoblotting.

For dot blot detection of total cellular lysates, cells were lysed in lysis buffer as above followed by denaturation in sample buffer, and were loaded onto nitrocellulose membrane (Bio-Rad) in a dot blot apparatus (GE Healthcare). Thereafter, the membrane was dried for standard immunoblotting.

### Confocal fluorescence light microscope image acquisition and analysis

The following antibodies were used for immunostaining: anti-myc (Millipore); anti-MAP1LC3B (Novusbio); anti-Ubiquitin (Santa Cruz); anti-p62 (Enzo); anti-polyQ (Sigma), goat anti-rabbit Alexa Fluor 555 (Thermo), and goat anti-mouse CF350 (Sigma).

Cells were seeded onto glass coverslips in 6-well cell culture plates. Cells were washed with PBS and fixed in 4% paraformaldehyde (Sigma) for 10 min, permeabilized with 0.1% Triton X-100 (Sigma) for 5 min, and blocked in 5% milk in PBS at room temperature for 1 hour. Primary antibodies were applied at a dilution of 1:50-1:100 in block overnight at 4°C, then washed in PBS (2 x 5 min) and incubated with secondary antibodies at a dilution of 1:1000 in dark at room temperature for 1 hour. Coverslips were washed with PBS (3 x 5 min), mounted onto glass slides with Fluoromount (Sigma).

For staining with ER-Tracker Red or LysoTracker Red DND-99 (Invitrogen), the dyes were added to the culture prior to fixation or vitrification according to the manufacturer’s protocol. 30 min prior to vitrification or live confocal light microscope imaging, cells were also stained with Annexin V-pacific blue (Invitrogen) or iFlour 555 (Abcam) according to the manufacturer’s protocol to quantify and exclude apoptotic cells from further analysis, respectively.

Fluorescence microscopy data were acquired at the MPI imaging facility, on a confocal laser scanning microscope (Zeiss LSM 780, Jena, Germany) with a PLAN/APO 63x oil objective and the Zeiss Zen software. Images acquired with the Airyscan 2 detector was performed on the Zeiss LSM 980 microscope. For fixed slides, images for quantifications of control and the treatments were taken with the same exposure setting at around the aggregate equator, data analysis was performed in Fiji (Schindelin et al., 2012). Representative datapoints from reproducible experiments shown. The measurements of the aggregate cross-sectional areas were carried out with the central zone thresholded to > 8000 for GFP intensity (threshold tool), and the peripheral region set to 800-8000 for the intensity threshold. Fluorescence analyses of LysoTracker and MAP1LC3B staining were performed similarly using threshold tool with signal intensity >800 across samples, the area within ∼1.5 μm around the central zone was quantified quantified. Data analyses were performed in GraphPad Prism 9 (Graphstats Technologies), p values generated from two-tailed student’s paired t-test.

Confocal live imaging was acquired in Zeiss LSM 980 Airyscan 2 with PLAN/APO 63x oil objective, with cells in 5 % CO_2_ and 37 °C. Z-stacks of the aggregates were generated by scanning at 0.5 µm interval for a 7 µm span. FRAP assay was carried out for the tracking with single frames imaged at 20 s intervals for ∼400s. FRAP analysis was performed with double normalization using the easyFRAP software tool (Koulouras et al., 2018).

### Isolation of mCherry-LC3B and 97Q-GFP vesicles and LC-MS/MS

The isolation of mCherry-LC3B and Htt97Q-GFP vesicles was carried out as following (Gao et al., 2010) with minor modifications. Co-transfected HEK293 LC3B knock out or Neuro2a were grown in 6-well plates for 48 hours with or without treh/rapa. For chloroquine treatment, 100μM was applied for 5 hours prior to harvest. Cells were harvested and homogenized with 22-gauge needle in 1.5ml cold isotonic buffer (0.25M sucrose, 1mM EDTA, 20mM HEPES pH 7.4) with protease inhibitor. Lysates were centrifuged at 800g at 4°C for 10 min to pellet the nuclei. The supernatant was then centrifuged at 10,000g for 20 min at 4°C, the pellet was collected and washed in PBS to remove free mCherry-LC3B. The pellet was then resuspended in PBS and filtered (Corning) to remove clumps prior to vitrification, analyses by confocal microscope or by flow cytometry sorting (Imaging facility).

Flow cytometry sorting was performed with a 70 µm nozzle on a BD FACSAria III with standard calibrations by fluorescence beads. A negative control without the tagged proteins was included as background control. Vesicles < ∼1µm diameter were sorted and collected in PBS, and were applied to dot blots or processed for LC-MS/MS (Biochemistry core facility) in triplicates.

For LC-MS/MS, proteins were reduced and alkylated in SDC buffer (1% Sodium deoxycholate, 40 mM 2-Cloroacetamide (Sigma), 10 mM TCEP (Thermo) in 100 mM Tris pH 8.0) for 20 min at 37 °C. The samples were then diluted with MS grade water (VWR) and digested overnight at 37 °C with 1 µg Lys-C (Labchem-wako) and 2µg trypsin (Promega), followed by acidification with Trifluoroacetic acid (Merck) to 1% (pH < 2). Next, the samples were purified via Sep-Pak Vac 1cc (50mg) tC18 Cartridges (Waters GmbH) with 0.1M acetic acid (Roth) wash and eluted with 80% Acetonitrile and 20mM acetic acid (Roth), and vacuum dried for resuspension in Buffer A (0.1% (v/v) Formic acid (Roth)). Peptides were then loaded onto a 30-cm column (packed with ReproSil-Pur C18-AQ 1.9-micron beads, Dr. Maisch GmbH) via the Thermo Easy-nLC 1200 autosampler (Thermo) at 60°C. Using the nano-electrospray interface, peptides were sprayed onto the Orbitrap MS Q Exactive HF (Thermo) in buffer A at 250 nl/min, and buffer B (80% Acetonitril, 0.1% Formic acid) was ramped to 30% in 60 min, 60% in 15 min, 95% in 5 min, and finally maintained at 95% for 5 min. MS was operated in a data-dependent mode with survey scans 300 - 1650 m/z (resolution of 60000 at m/z =200), and up to 10 top precursors were selected and fragmented using higher energy collisional dissociation (HCD with a normalized collision energy value at 28). The MS2 spectra were recorded at a resolution of 30000 (at m/z = 200). AGC target for MS and MS2 scans were set to 3E6 and 1E5 respectively within a maximum injection time of 100 and 60 ms for MS and MS2 scans respectively. Dynamic exclusion was set to 30ms.

Raw MS data were processed using the MaxQuant platform (Cox and Mann, 2008) with standard settings, and searched against the reviewed Human or Mouse Uniprot databases, as well as mCherry-LC3B and polyQ-GFP sequences, allowing precursor mass deviation of 4.5 ppm and fragment mass deviation of 20 ppm. MaxQuant by default enables individual peptide mass tolerances. Cys carbamidomethylation was set as static, and Ub, Met oxidation and N-terminal acetylation as variable modifications. Protein abundances within a sample were calculated using iBAQ intensities (Schwanhausser et al., 2011) and were quantified over the samples using the LFQ algorithm (Cox et al., 2014) for analysis with the Perseus software (https://maxquant.net/perseus/).

### Anti-GFP pull-down with syringe filter partition

Htt64Q-GFP induced in Neuro2a for two days in a 15cm dish, wash with PBS, and were lysed in cold isotonic buffer (0.25M sucrose, 1mM EDTA, 20mM HEPES pH 7.4) with protease inhibitor using gauge 22-needle. Nuclei were pelleted at 800g for 10min at 4°C, supernatant was then centrifuged at 16000g for 10min at 4°C to pellet aggregates and the associated proteins. The pellet was washed once in PBS to remove soluble Htt64Q-GFP and was then dissolved in PBS, and passed through pre-equilibrated syringe filters with 0.2um or 0.4um pore sizes (Millipore). The filter blocked fractions were collected and pelleted again at 16,000g for 10min at 4°C. The fractions were then incubated with magnetic GFP-Trap (ChromTek) for 6 hours at 4°C in binding buffer (140 mM NaCl, 25 mM Tris pH 7.6–8.0, 0.5% NP40, 1 mM EDTA) and 2% BSA (Sigma). Lysates from cells without muristerone A induction was used as negative control in pull down. After 5 washes with the binding buffer, the bound fraction was eluted and denaturation in sample buffer for dot blot analysis.

### Sample vitrification for cryo-ET

Cells were seeded on holey carbon-coated 200 mesh gold EM grids (Quantifoil Micro Tools, Jena, Germany) in 35 mm cell culture dishes. Prior to vitrification, cells were applied with DMEM containing 10 % glycerol as a cryo-protectant and Dynabeads (Invitrogen) at 1:40 dilution for 3D CLEM workflow, then immediately mounted on Vitrobot Mark IV (Thermo), blotted from the back side using FEI Vitrobot Perforated Filter Paper (Whatman) with force 10 for 15 seconds at room temperature, and plunged into a 2:1 ethane:propane mixture cooled down to liquid nitrogen temperature. Plunge-frozen grids were then clipped into Autogrid support frames modified with a cut-out (Thermo), stored in liquid nitrogen, and maintained at ≤−170°C for all steps.

For the vitrification of isolated vesicles, 4 μl of the sample was applied to glow discharged holey carbon-coated 200 mesh copper EM grids (Quantifoil Micro Tools), vitrified as above at 4°C and 100 % humidity.

### Correlated light-electron microscopy (CLEM) and cryo-focused ion beam (FIB) milling

Vitrified sample on autogrids were loaded onto a cryo-confocal LM set up (Leica SP8) equipped with a 50X/0.9 NA objective (Leica Objective), metal halide light source (EL6000), air-cooled detector (DFC900GT), a cryo-stage (-195 °C), and two HyD detectors. The sample was kept in liquid nitrogen vapor, following a similar workflow as described (Schorb et al., 2017). Cryo-confocal z-stacks (step size 500 nm, x-y pixel size 85 nm) were taken with pin hole = 1, and a 9 μm depth with the LAS X Navigator software, using 488 and 552 nm laser excitation for GFP and RFP tagged proteins, respectively, also picking up signals from the auto-fluorescent Dynabeads. To improve signal clarity, image stacks were de-convoluted and restored with Huygens Essential software (Scientific Volume Imaging) to remove noise. The stack was then imported into the 3D correlation software (Arnold et al., 2016) and re-sliced into cubic voxels.

For preparing the lamella (150-250 nm), autogrids were mounted into a Quanta dual-beam 3D FIB/scanning electron microscope (Thermo) equipped with a transfer shuttle system (PP3000T, Quorum) at < −180 °C throughout milling. To protect the milling front of the lamella, gaseous organometallic platinum was sprayed onto the sample on the cryo-stage using a gas injection system. To target the cell for milling, the grid square was correlated with the cryo-confocal fluorescence z-stack using the 3Dcorrelation software. The target was imaged and correlated iteratively with the z-stack throughout milling for accuracy. The 12-15 μm wide lamellas were generated using a Gallium FIB at 30 kV with a 20° stage angle in three consecutive steps. The more distant region (>2 μm) above and below the target was rough milled with a higher current of 500 pA, followed by fine milling to a ∼800 nm lamella using a current of 100 pA. A final polishing of the lamella to thickness of 150-250 nm was carried out with a 30-50 pA current. Lamella final thickness was estimated with SEM at 3 keV, for a lack of overcharging.

### Cryo-ET data acquisition, tomogram reconstruction, and analysis

Lamellas were imaged in a FEI G2 Polara or Titan Krios cryo-TEM equipped with a field emission gun operating at 300 kV, a post-column energy filter (Gatan, Pleasanton, CA, USA) operating at zero-loss, and a 4k x 4k K2 Summit direct electron detector (Gatan). The energy filter was used to increase image contrast with a slit width of 20 eV. Low-magnification were taken at 5600x (object pixel size 2.18 nm) to generate lamella overviews. High-magnification (18,000x, 27,500x, 34,000x, 42,000x) with a pixel size of 0.65, 0.42, 0.34, 0.32 nm respectively) tilt series were recorded at sites of interest using the SerialEM software (Mastronarde, 2005), operating in low dose mode with tracking and focus enabled. Tilt series were taken with a 2° tilt increment with an angular range from ∼−60° to 60°. The K2 camera operating in dose fractionation counting mode, recorded frames every 0.2 s for ∼2 electrons/A² per tilt angle. For the tilt series, the cumulative dose was in the range of 90-120 electrons/Å². Targeted defocus of -9 μm (Polara) or -5 μm (Krios) were applied to boost contrast. The low magnification views (5600x) of the lamella provided enough detail to locate the aggregates and surrounding structures for tomogram acquisitions.

For tilt series acquisition of the isolated vesicles, Htt97Q-GFP and LC3B-RFP positive puncta were first located on the vitrified grid using the Leica cryo-confocal LM. Sample features including holes and cracks were used as landmarks for target identification.

K2 camera raw frames were preprocessed using in-house Matlab (Nickell et al., 2005) wrapper scripts (Tomoman: https://github.com/williamnwan/TOMOMAN). The relative shifts of the image between camera frames due to stage drift and beam-induced motion were corrected by MotionCor2 (Zheng et al., 2017), followed by exposure filtering (Grant and Grigorieff, 2015). The tilt series were then aligned using patch tracking, binned by 4, and reconstructed by weighted back projection in IMOD (Kremer et al., 1996). FIB-related imperfections of the lamella were removed for image display, using the LisC filter algorithm (Felix J.B. Bäuerlein, 2022). Tomogram contrast was improved using Topaz (Bepler et al., 2020) or cryo-care (Buchholz et al., 2019). Ice contamination were removed from the tomogram using the MaskTomRec software (Fernandez et al., 2016). The overlays of fluorescence z-stack, SEM, and TEM images were generated using the Transform/Landmark correspondence plugin (Fiji).

Tomogram segmentation was performed in Amira (Thermo). Membranes were automatically detected by TomoSegMemTV using tensor voting (Martinez-Sanchez et al., 2014), followed by manual refinement in Amira. polyQ fibrils were detected using the XTracing module (Rigort et al., 2012). In brief, tomograms were denoised by a non-local means filter, and the fibril containing regions were searched for a cylindrical template of 8 nm (diameter) and 42 nm (length). The resulting cross-correlation fields were adjusted to a range of 0.68-0.8 for optimized detection. The amorphous aggregate density was approximated by fluorescence correlation, the corresponding region was segmented by the magic wand tool in Amira. The cytosolic ribosomes were detected by PyTom template matching (Hrabe et al., 2012), using a low pass filtered (60Å) ribosome template (EMDB: 5592).

From the ∼300 aligned and exposure filtered tomograms (Polara) without additional contrast enhancement, the diameters and relative intensities of the phagophores and autophagosomes, as well as the downstream vesicles (autolysosomes and lysosomes) were quantified in Fiji. Diameters (nm) were taken as the longest distance of the inner bi-lamellar membrane of phagophores and autophagosomes, or the longest distance between the uni-lamellar downstream vesicles. Relative content intensity was calculated as the average intensity of the volume inside the phagophores and autophagosomes, or the downstream vesicles, normalized to the average intensity of the entire tomogram, excluding regions with ice crystals and broken edges. p values generated from two-tailed student’s paired t-test.

---

**Figure S1.**
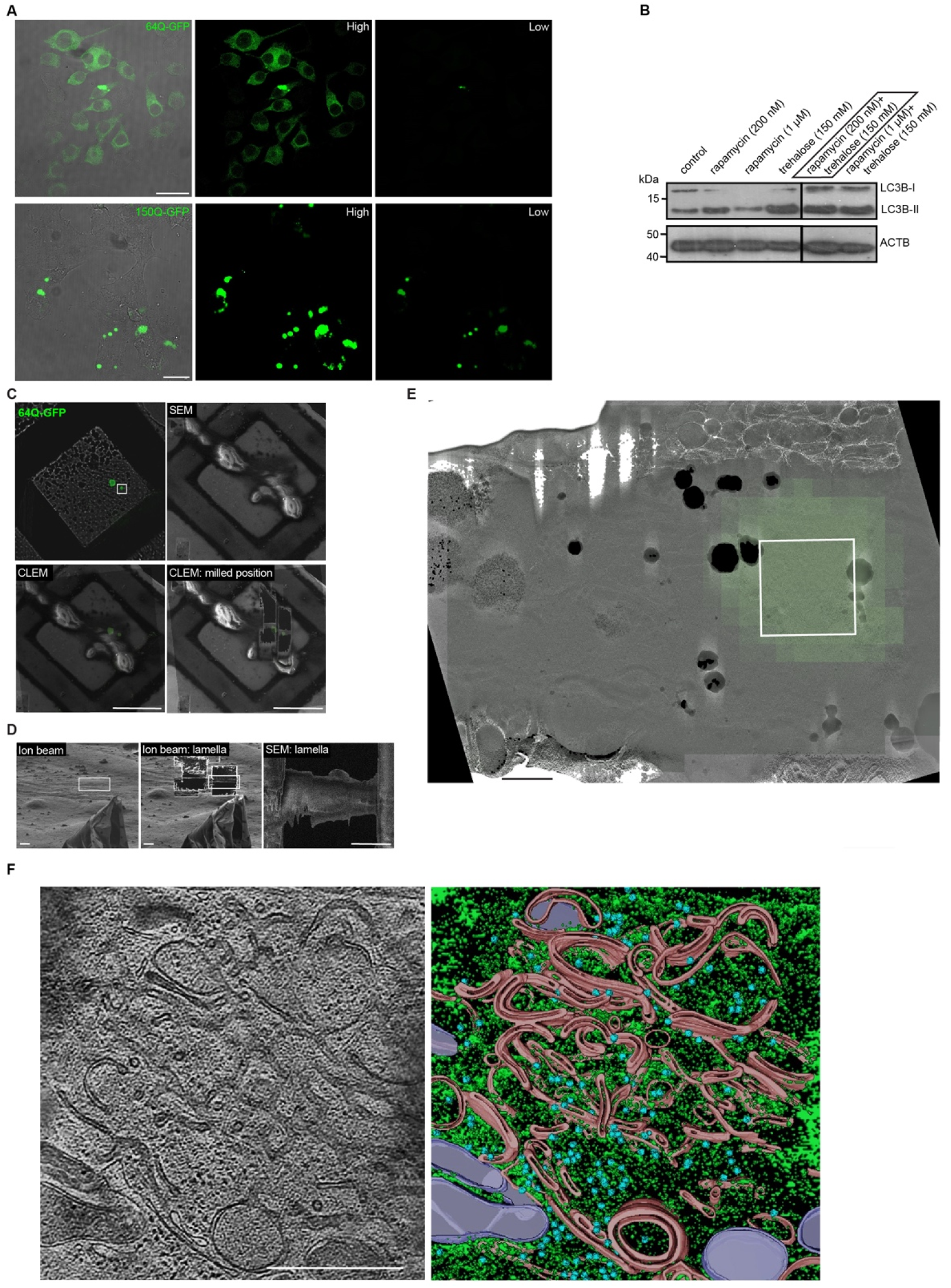
PolyQ aggregates with varying repeats are differentially targeted by autophagy (related to Figure 1) **(A**) Expression patterns of 64Q- and 150Q-GFP in Neuro2a upon 2-day muristerone A induction, displayed with high (H) and low (L) exposures for the fluorescence channel. **(B)** Autophagy induction in Neuro2a immunoblotted against LC3B, ACTB as loading control, boxed condition used for all assays. **(C)** cryo-CLEM workflow: Neuro2a expressing 64Q-GFP grown on a cryo-EM grid, imaged in cryo-confocal for z-stacks for correlation during targeted focused ion beam (FIB) milling. Correlation with the SEM images shown before and after milling. Site of interest boxed in white. **(D)** FIB views of the target before and after milling; SEM lamella image. **(E)** TEM Lamella view (5600x, LisC-filtered) indicating the tomogram area, overlaid with fluorescence signal. **(F and G**) Tomographic slice of a dim 64Q-GFP aggregate as a mixture of amorphous content with curved membrane structures resembling phagophores (PH) and autophagosomes (AP). Segmentation (G) with PH, AP, and ER related membranes (pink), mitochondria (purple), ribosomes (blue), amorphous polyQ (green). Scale bars: 20μm in (A), 50μm in (C), 5μm (D), 1μm in (E), 500nm in (F, G).

**Figure S2.**
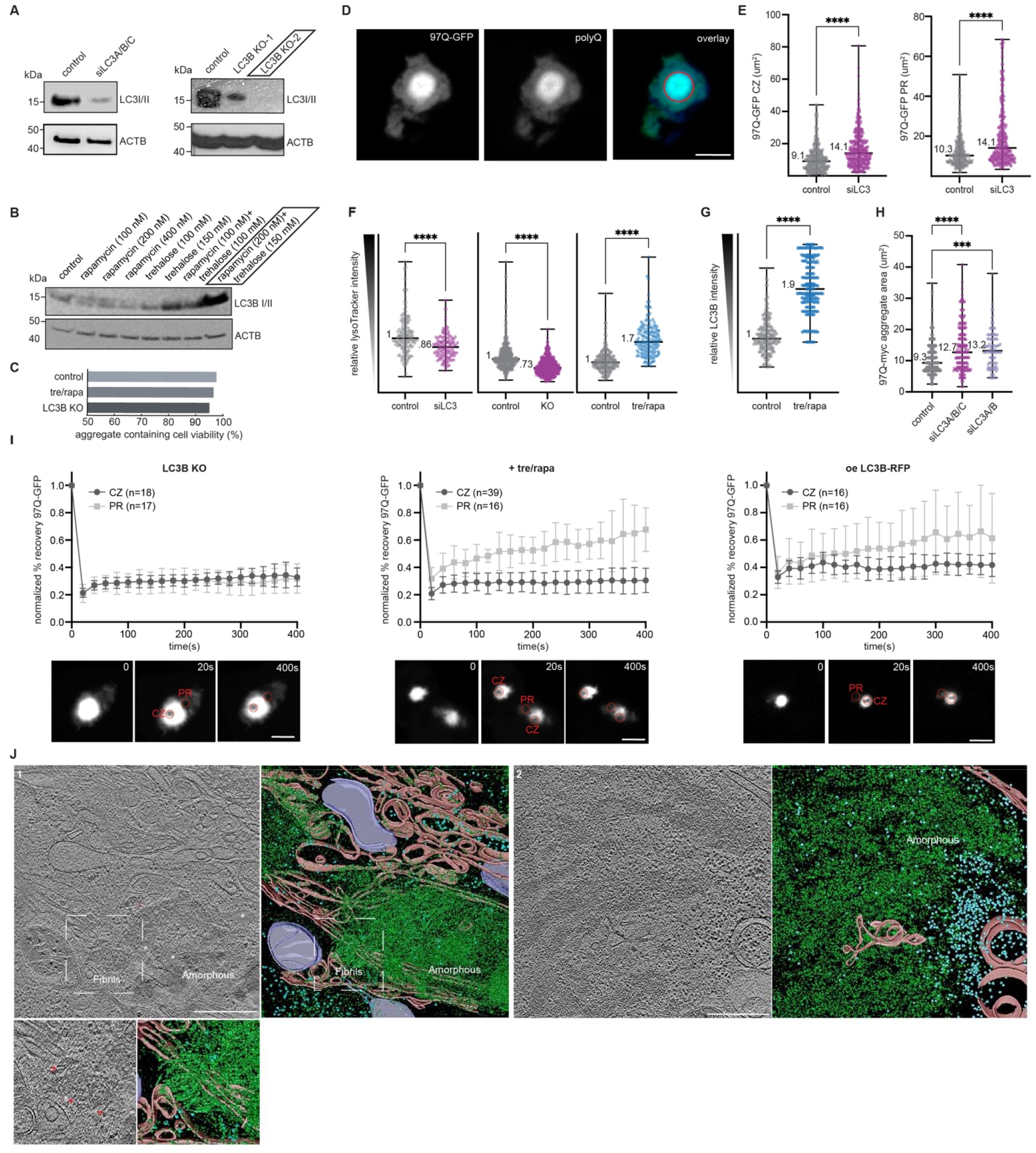
Autophagy engages the different phases of 97Q aggregates (related to Figure 2) **(A**) Validation of the LC3 siRNA knock down, or CRISPR/Cas9 LC3B knock out (KO) by immunoblotting against LC3B, ACTB as loading control (HEK293). LC3B KO2 was used for all assays. **(B**) Autophagy induction in HEK293 by immunoblotting against LC3B, ACTB as loading control, boxed condition was used for all assays. **(C**) Fluorescence live imaging for cell viability upon Annexin V-iFlor 555 staining, control n=272, treh/rapa n=322, and LC3B knock out n=492. **(D**) Representative confocal images of 97Q-GFP in LC3B knock out, co-stained with antibodies against polyQ, red circles approximate central zone, the rest as peripheral region. **(E**) Image quantification of 97Q-GFP in control or upon LC3 siRNA knock down for central zone (CZ: control n=323, siLC3 n=361) and peripheral region (PR: control n=265, siLC3 n=305) with medians displayed, CZ and PR intensities thresholded to >8000 and 800-8000, respectively, ****p<0.0001. **(F**) Image quantification of LysoTracker relative intensity (area x signal intensity) within 1.5μm around the central zone in control v. siLC3A/B/C (control n=153, knock down n=136); v. LC3B knock out (control n=402, KO n=551); and v. treh/rapa (control n=217, +treh/rapa n=158) with medians displayed, ****p<0.0001. **(G**) Image quantification of LC3B relative intensity within 1.5μm around the central zone in control v. treh/rapa (control n=209, +treh/rapa n=334) with medians displayed, ****p<0.0001. **(H**) Image quantification of 97Q-myc aggregate area: control n=337, siLC3A/B/C n=232, siLC3A/B n=111 with median values displayed, ****p<0.0001. **(I**) FRAP analysis with double normalization in the EasyFrap software, for the 97Q-GFP central zone (CZ) and peripheral region (PR) in HEK293 upon LC3B knock out (left), treh/rapa induced autophagy (middle), or mCherry-LC3B co-expression (right). Arrows indicate the time point of bleaching, assayed for ∼400s at 20s intervals; shaded regions indicate s.d. Representative images below indicate bleach sites in central and peripheral regions (red circles). **(J**) Tomographic slices of polyQ phase as an amorphous-fibrillar mixture (1) with enlarged panel below for the boxed fibrillar region, contamination ice (marked by *) cleaned by MaskTomRec; or as a loose amorphous phase (2) separated from ribosomes and organelles. Segmentation: ER related membranes (pink), mitochondria (purple), ribosomes (blue), and polyQ (green). [CLEM workflow and lamella view: SI Figure 1D-F] Scale bars: 5μm in (D), 5μm in (I), 500nm in (J).

**Figure S3.**
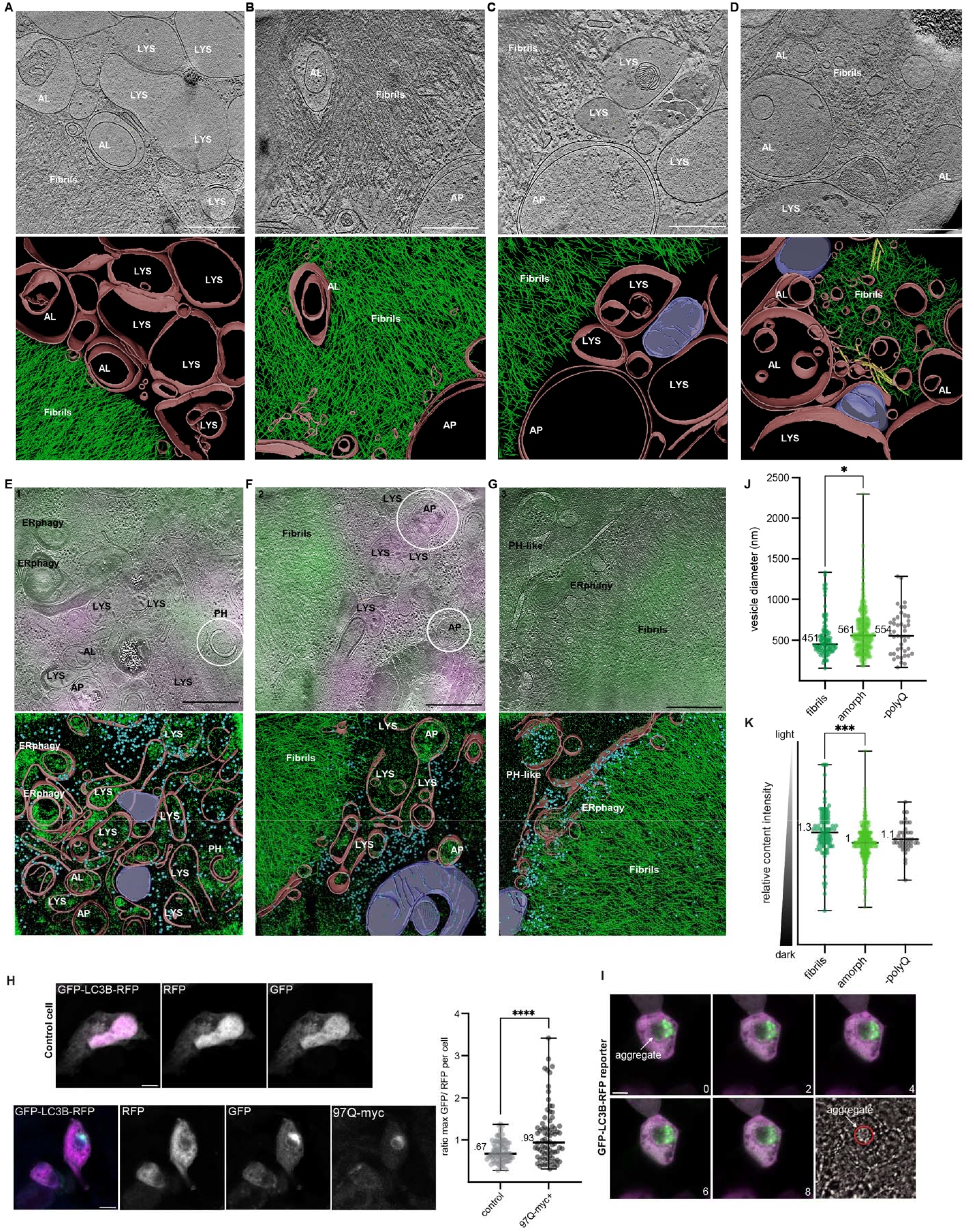
Autophagic structures prefer amorphous over fibrillar cargo *in situ* (related to Figure 3) **(A-D**) Tomographic slices with 97Q-GFP aggregates and LysoTracker upon induced autophagy. Autophagosomes (AP), autolysosomes (AL), and lysosomes (LYS) recruited to the fibrils are empty. [CLEM and lamella views: A (SI figure 5A-C), B-C (A (SI 5D-F), D (A (SI 5G-I)]. **(E-G)** Tomographic slices with 97Q-GFP aggregates and mCherry-LC3B superimposed with fluorescence, AP and phagophore (PH) are circled. E is the full tomographic slice for Figure 3E. [CLEM and lamella views: SI figure 4I-K]. Segmentation with PH, AP, AL, LYS, and ER related membranes (pink), mitochondria (purple), ribosomes (blue), microtubule (yellow), and polyQ (green). **(H and I**) Co-expression of GFP-LC3B-RFP reporter and 97Q-myc (blue), max GFP to RFP per cell quantified to approximate aggregate effect on autophagy flux, with medians displayed (H). Live imaging of the GFP-LC3B-RFP reporter with 97Q-myc viewed in DIC channel, showing the same z-slice at 2 min interval for 8 min (I). **(J and K)** Tomogram quantification of the downstream autophagic structures (autolysosomes, lysosomes): diameter (J) and content density (K) proximal to fibrils (n=93), to amorphous polyQ (n = 398), or in cells without polyQ (n = 43) with medians displayed, ***p=0.001, *p=0.05. Scale bars: 500nm in (A-G), 5μm in (H-I).

**Figure S4.**
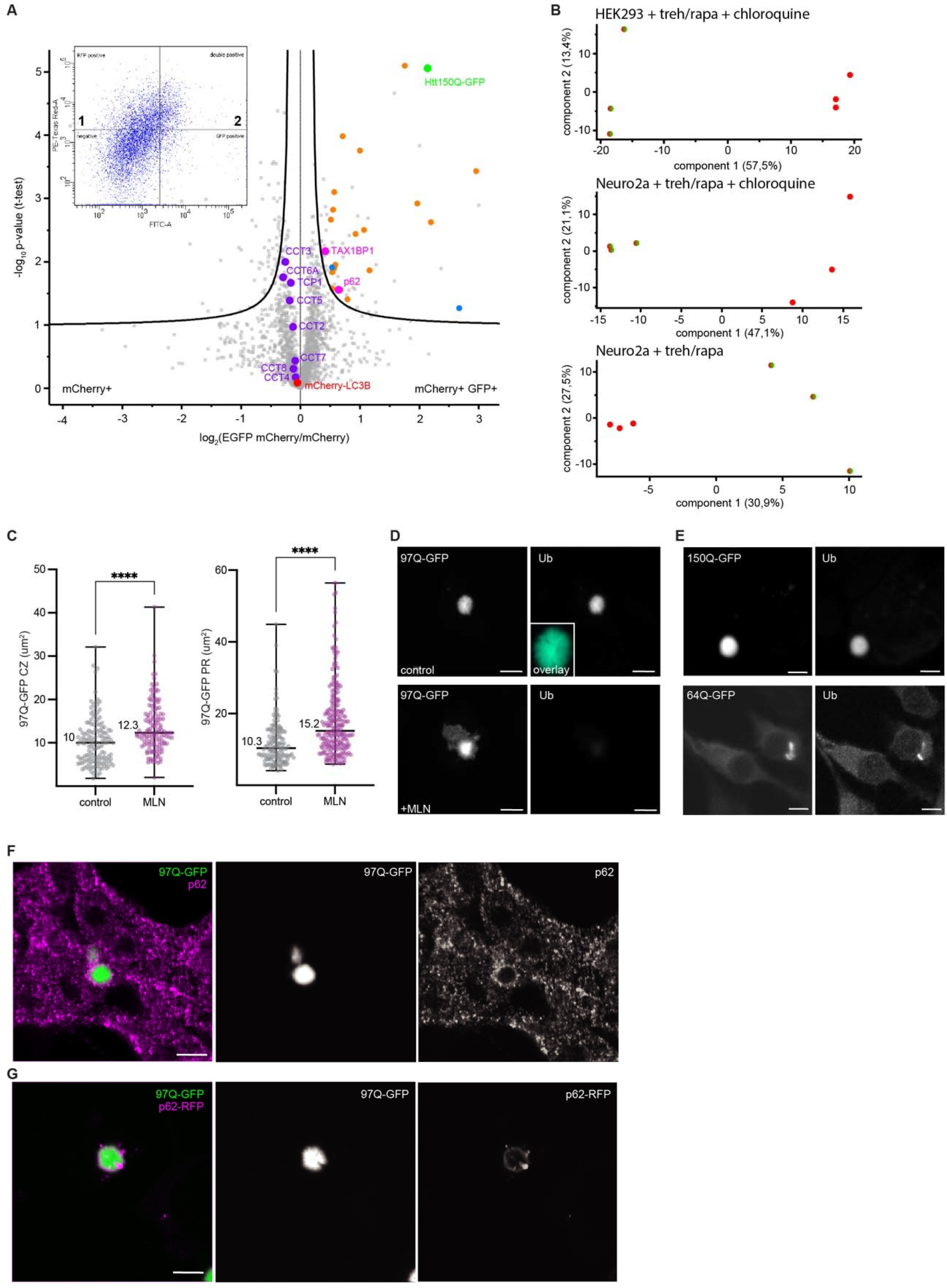
The autophagic intake of the polyQ is a p62 mediated process (related to Figure 4) **(A)** Label-free quantitative mass spectrometry analysis of the sorted vesicles with mCherry+ controls v. mCherry+ GFP+ from Neuro2a + treh/rapa. Insert: vesicles sorting for mass spectrometry with mCherry+ (quadrant 1) and mCherry+ GFP+ (quadrant 2). Receptors for Ub-substrates (pink), TRiC subunits (purple), and RNA processing factors (blue) are highlighted; proteasome and protein quality control regulators (orange) and are colored. **(B)** Mass spectrometry principle component analyses for mCherry+ (red) v. mCherry+ GFP + (orange) in triplicates. **(C)** Image quantification of 97Q-GFP central zone (control n=140 v. +MLN7243 n=136), and the peripheral region (control n=140 v. +MLN7243 n=227) with medians displayed, ****p<0.0001. **(D)** Representative confocal images of 97Q-GFP control or with MLN7243 (0.2 µM, 12 hours). **(E)** Representative confocal images of 150Q- and 64Q-GFP in Neuro2a, stained with antibody against Ub. **(F and G)** Representative confocal images of 97Q-GFP stained with p62 (F) or co-expressed with p62-RFP (G) in HEK293. Scale bars: 5μm in (D-G).

**Figure S5.**
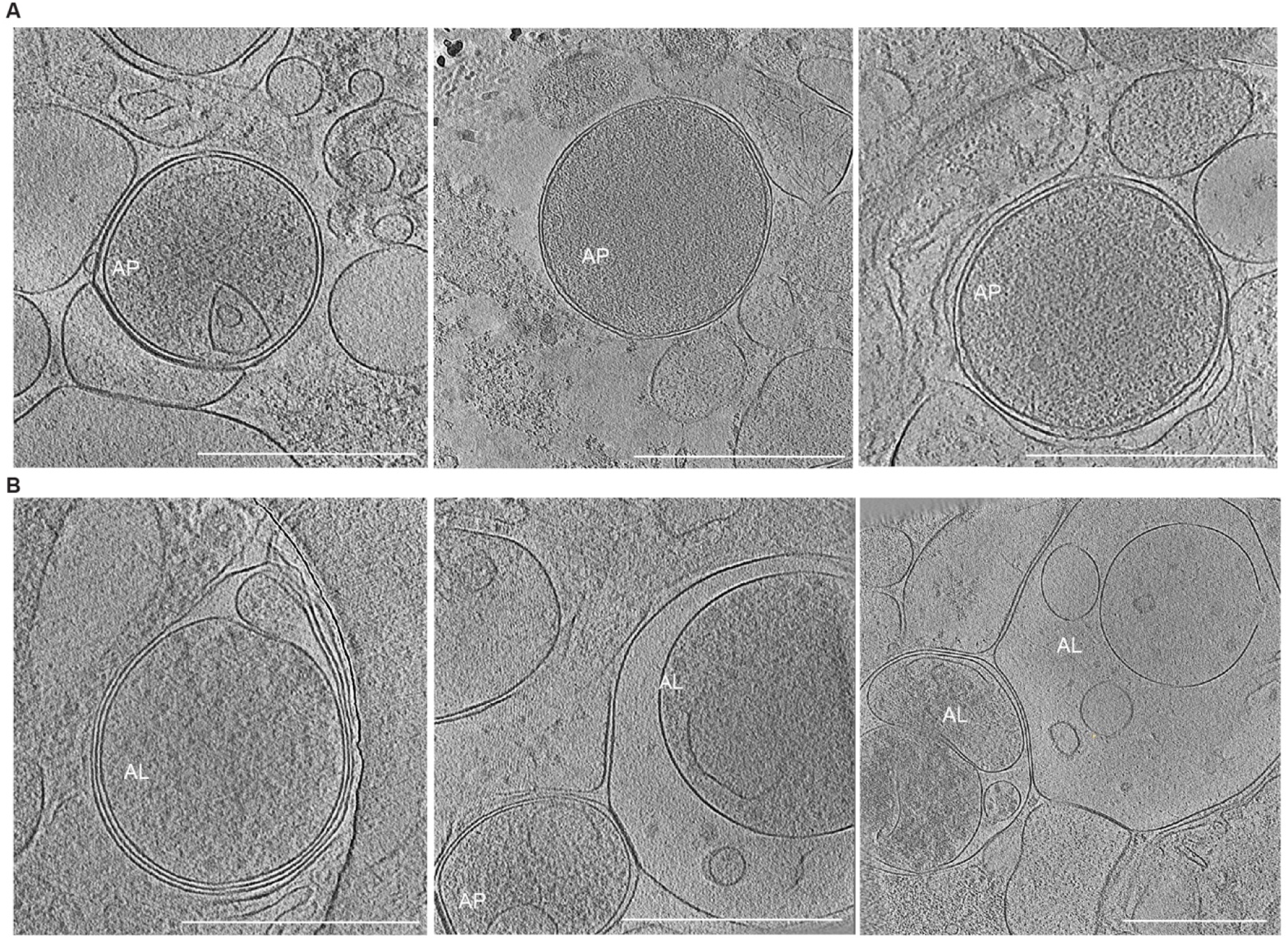
Autophagy preferentially intakes the amorphous phased 97Q (related to Figure 5) (A and B) Additional tomographic slices (27,500x-34,000x) of 97Q-GFP and mCherry-LC3B positive autophagosomes (A) and autolysosomes (B). Scale bars: 500nm.

**SI Figure 1.**
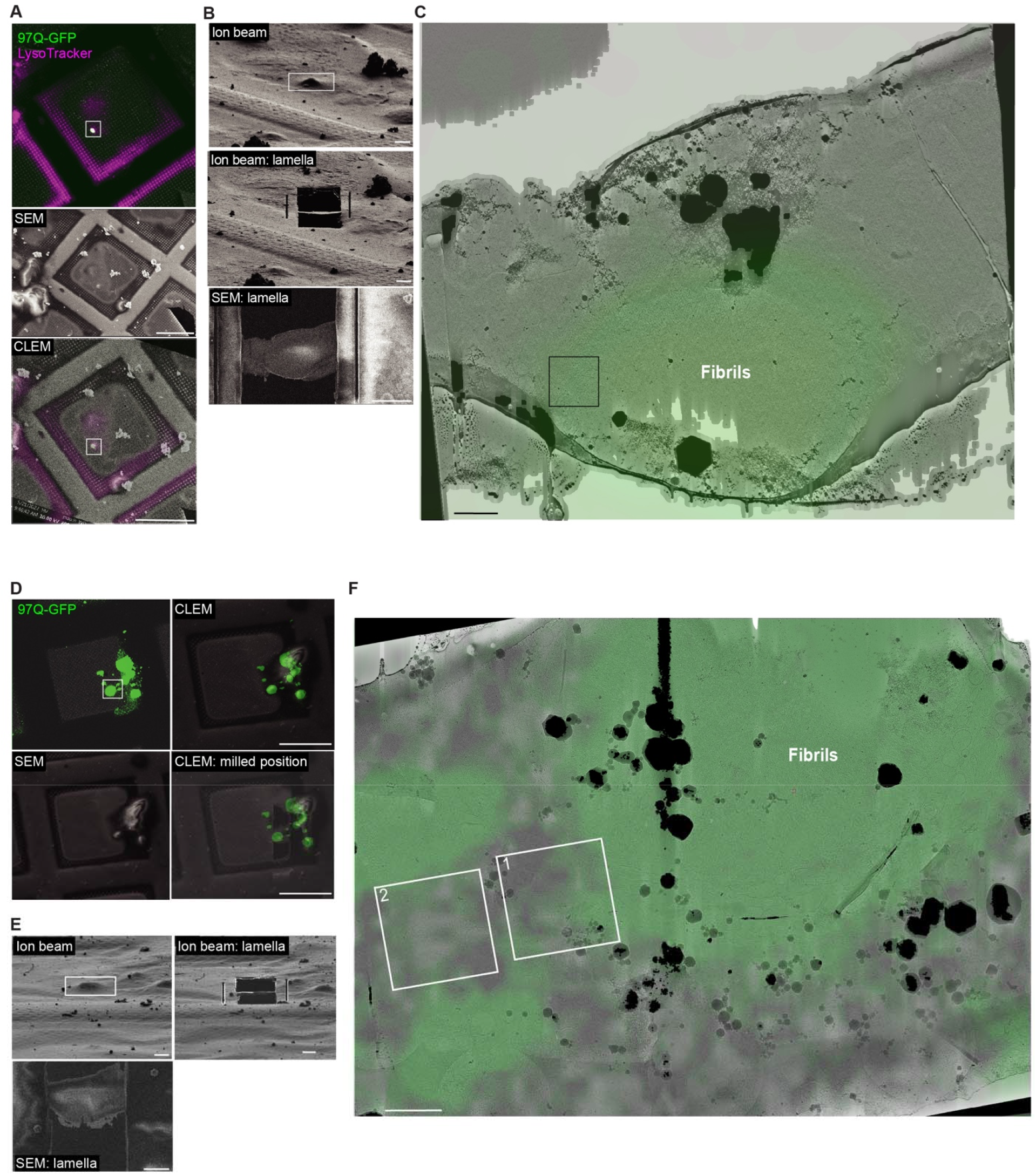
The amorphous polyQ phase *in situ* by cryo-ET (related to Figure 2) Related to Figure 2F-G (A) Cryo-CLEM workflow: 97Q-GFP stained with LysoTracker, site of interest boxed. (B) FIB and SEM views of the target, site of interest boxed. (C) Lamella view (5600x) indicating the tomogram area, overlaid with GFP signal. Scale bars: 50 μm in (A), 5 μm in (B) 1 μm in (C). Related to Figure S2J (D) Cryo-CLEM workflow: LC3B knock out cell expressing 97Q-GFP, site of interest boxed. (E) FIB and SEM views of the target, site of interest boxed. (F) Lamella view (5600x) with the tomogram areas indicated, overlaid with GFP signal. Scale bars: 50 μm in (D), 5 μm in (E) 1 μm in (F).

**SI Figure 2.**
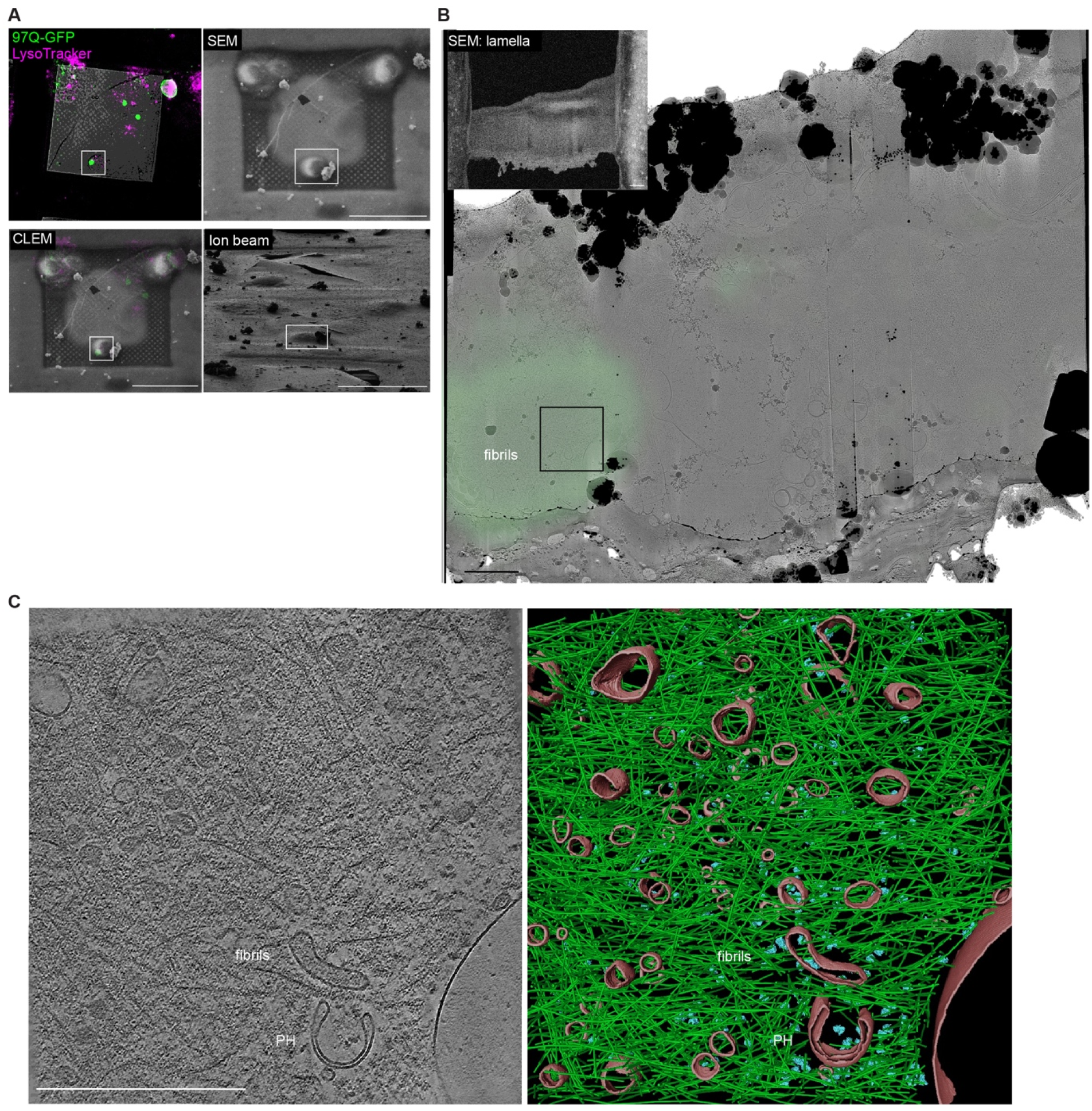
Empty phagophore trapped by fibrils (related to Figure 3B) (A) Cryo-CLEM workflow: 97Q-GFP stained with LysoTracker, site of interest boxed. (B) SEM and TEM Lamella view (5600x) indicating the tomogram area, overlaid with GFP signal. (C) Tomographic slice (42000x). Segmentation with phagophore (PH) and ER related membranes (pink), ribosomes (blue), and polyQ (green). Scale bars: 50 μm in (A), 1 μm in (B), 500 nm in (C).

**SI Figure 3.**
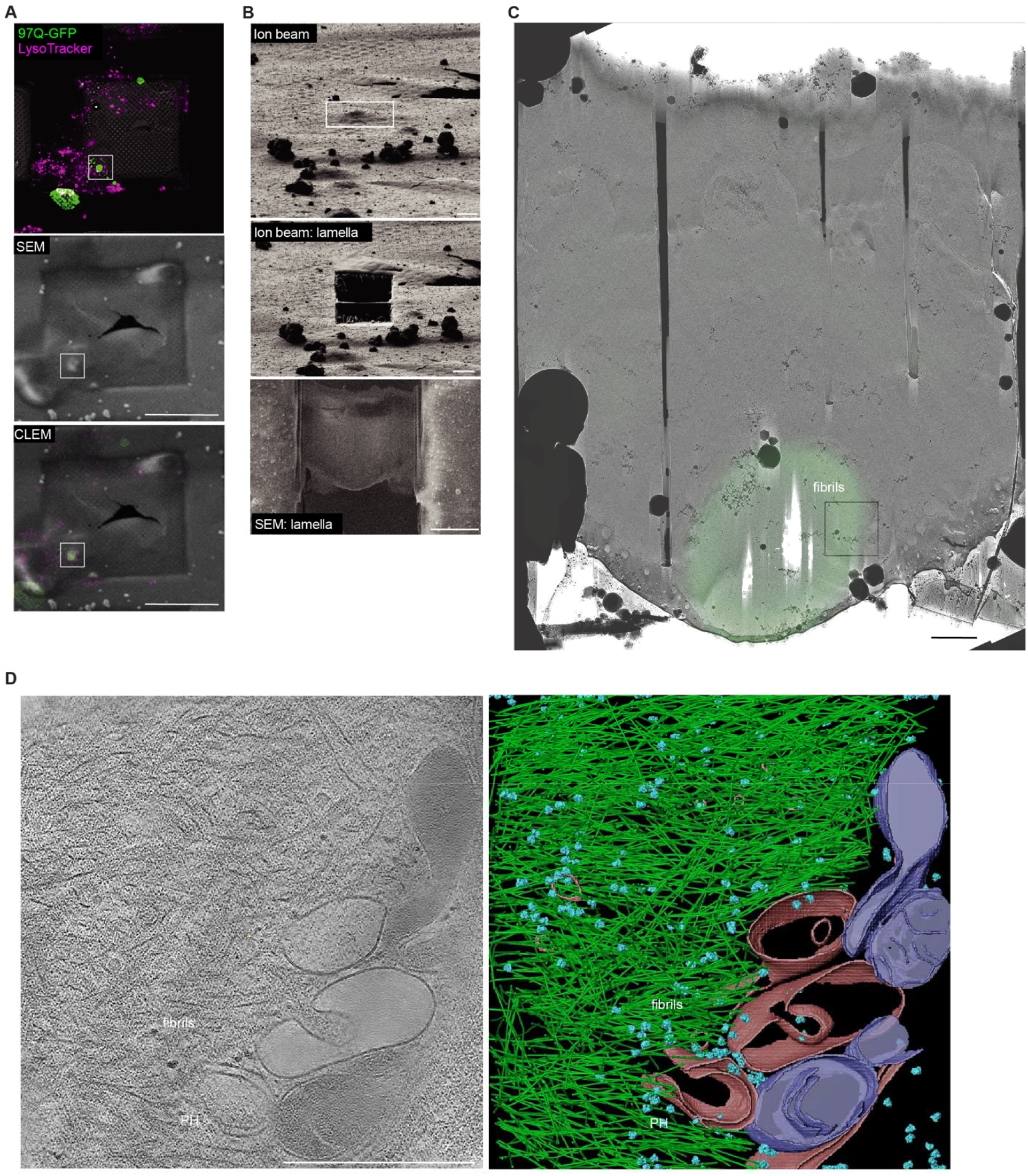
Empty phagophore trapped by fibrils (related to Figure 3C) (A) Cryo-CLEM workflow: 97Q-GFP stained with LysoTracker, site of interest boxed. (B) FIB and SEM views of the target, site of interest boxed. (C) Lamella view (5600x) indicating the tomogram area, overlaid with GFP signal. (D) Tomographic slice (42000x). segmentation with phagophore (PH) and ER related membranes (pink), mitochondria (purple), ribosomes (blue), and polyQ (green). Scale bars: 50 μm in (A), 5 μm in (B), 1 μm in (C), 500 nm in (D).

**SI Figure 4.**
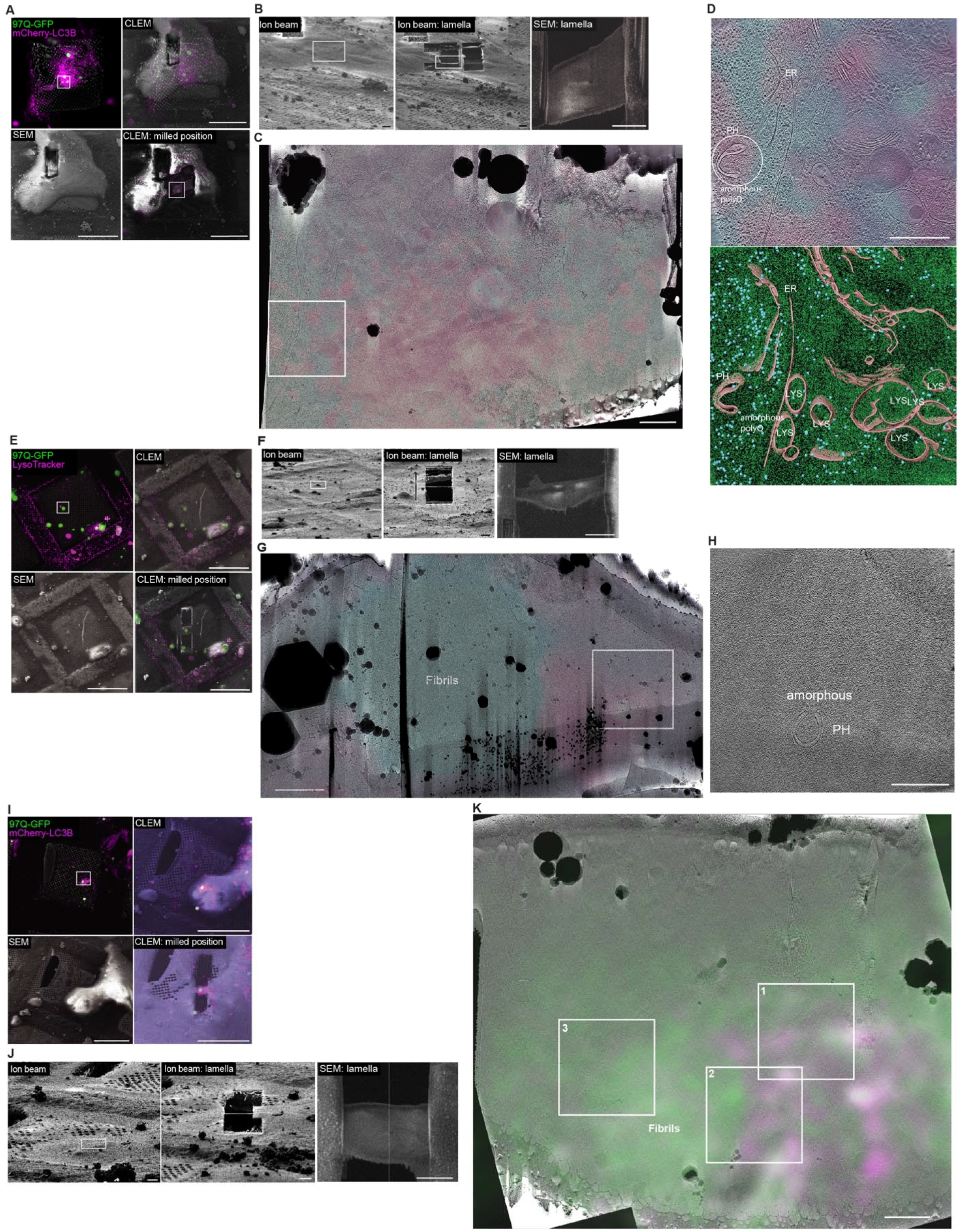
Phagophores with amorphous polyQ intake *in situ* (related to Figure 3) (A-D) An additional example of a phagophore trapped within the amorphous polyQ phase (A) Cryo-CLEM workflow: 97Q-GFP and mCherry-LC3B co-expression, site of interest boxed. (B) FIB and SEM views of the target, site of interest boxed. (C) Lamella views (5600x) indicating the tomogram area, overlaid with fluorescence signal acquired on the lamella post TEM. (D) Tomographic slice (18000x) overlaid with fluorescence, with correlated phagophore circled white. Segmentation with phagophore (PH), lysosomes, and ER (pink), ribosome (blue), and polyQ (green). Scale bars: 50 μm in (A), 5 μm in (B), 1 μm in (C), 500 nm in (D). (E-H) Related to Figure 3D (E) Cryo-CLEM workflow: 97Q-GFP with LysoTracker staining, site of interest boxed. (F) FIB and SEM views of the target, site of interest boxed. (G) Lamella view (5600x) indicating the tomogram area, overlaid with fluorescence signal. (H) Tomographic slice (18000x). Scale bars: 50 μm in (E), 5 μm in (F), 1 μm in (G), 500 nm in (H). (I-K) Related to Figure 3E and S3E-G (I) Cryo-CLEM workflow: 97Q-GFP and mCherry-LC3B co-expression upon induced autophagy, site of interest boxed. (J) FIB and SEM views of the target, site of interest boxed. (K) Lamella view (5600x) indicating the tomogram areas, overlaid with fluorescence signal acquired on the lamella post TEM. Scale bars: 50 μm in (I), 5 μm in (J), 1 μm in (K).

**SI Figure 5.**
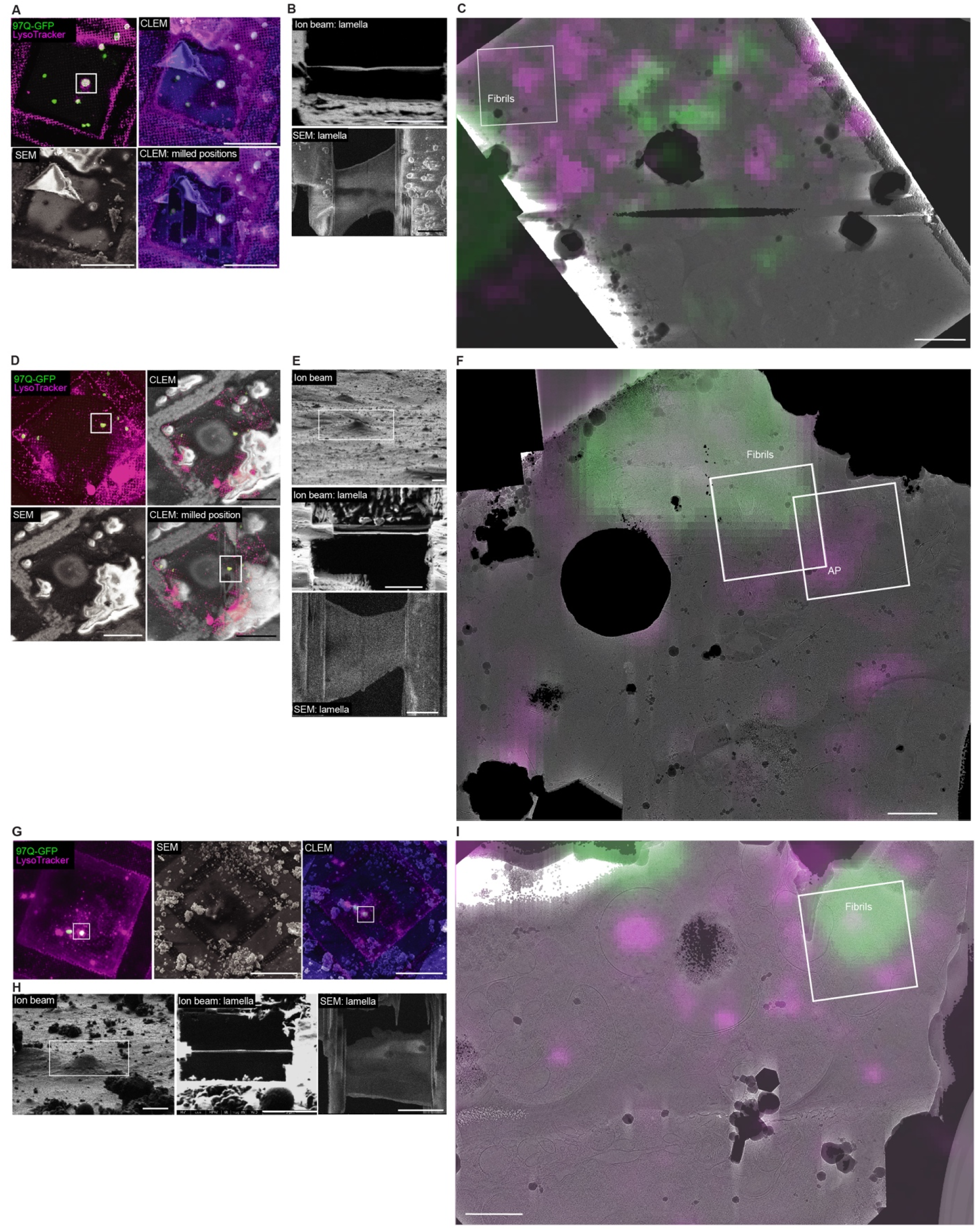
Downstream autophagic structures around the polyQ fibrils are empty (related to Figure 3) (A-C) Related to Figure S3A (A) Cryo-CLEM workflow: 97Q-GFP stained with LysoTracker, site of interest boxed. (B) FIB and SEM views of the target, site of interest boxed. (C) Lamella view (5600x) indicating the tomogram area, overlaid with fluorescence signal. Scale bars: 50 μm in (A), 5 μm in (B), 1 μm in (C). (D-F) Related to Figure S3B-C (D) Cryo-CLEM workflow: 97Q-GFP stained with LysoTracker, site of interest boxed. (E) FIB and SEM views of the target, site of interest boxed. (F) Lamella view (5600x) indicating tomogram areas, overlaid with fluorescence signal. Scale bars: 50 μm in (D), 5 μm in (E), 1 μm in (F). (G-I) Related to Figure S3D (G) Cryo-CLEM workflow: 97Q-GFP stained with LysoTracker, site of interest boxed. (H) FIB and SEM views of the target, site of interest boxed. (I) Lamella view (5600x) indicating the tomogram area, overlaid with fluorescence signal. Scale bars: 50 μm in (G), 5 μm in (H), 1 μm in (I).

**SI Figure 6.**
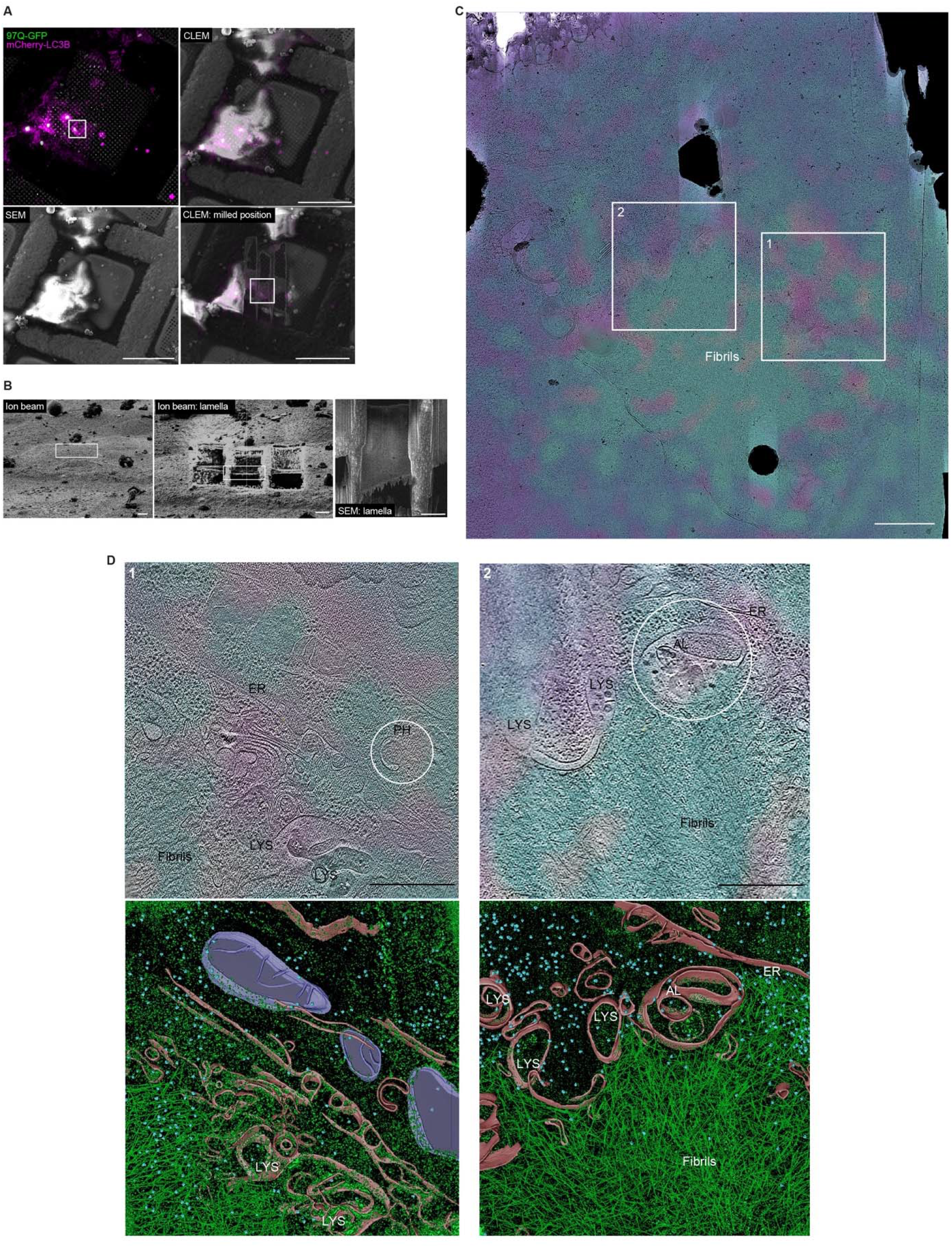
Phagophore and autolysosome with amorphous polyQ intake *in situ* (related to Figure 3) (A) Cryo-CLEM workflow: 97Q-GFP and mCherry-LC3B co-expression, site of interest boxed. (B) FIB and SEM views of the target, site of interest boxed. (C) Lamella view (5600x) indicating the tomogram areas, overlaid with fluorescence signal acquired in the cryo-confocal on the lamella post TEM. (D) Tomographic slice (18000x) overlaid with fluorescence, correlated phagophores (PH) and autolysosome (AL) circled in white, scale bars: 500nm. Segmentation with PH, AL, LYS, and ER related membranes (pink), ribosomes (blue), mitochondria (purple), and polyQ (green). Scale bars: 50 μm in (A), 5 μm in (B), 1 μm in (C), 500 nm in (D).

